# A patient-centric knowledge graph approach to prioritize mutants for selective anti-cancer targeting

**DOI:** 10.1101/2024.09.29.615658

**Authors:** Marina Gorostiola González, Adriaan P. IJzerman, Gerard J.P. van Westen

**Affiliations:** Leiden Academic Centre of Drug Research, Leiden University, 55 Einsteinweg, 2333 CC. Leiden, The Netherlands; Oncode Institute, the Netherlands

**Keywords:** Cancer, personalized oncology, mutation, selectivity, knowledge graph, network analysis, kinase, receptor tyrosine kinase

## Abstract

Personalized oncology has revolutionized cancer treatment by targeting specific genetic aberrations in tumors. However, the identification of suitable targets for anti-cancer therapies remains a challenge. In this study, we introduce a knowledge graph approach to prioritize cancer mutations with clinical, functional, and structural significance as potential therapeutic targets. Focusing on the human kinome, we integrate protein-protein interaction and patient-centric mutation networks to construct a comprehensive network enriched with transcriptomic, structural, and drug response data, together covering five layers of information. Moreover, we make the constructed knowledge graph publicly available, along with a plethora of scripts to facilitate further annotation and expansion of the network. Interactive visualization resources are also provided, ensuring accessibility for researchers regardless of computational expertise and enabling detailed analysis by cancer type and individual layers of information. This comprehensive resource has the potential to identify relevant mutations for targeted therapeutic interventions, thereby advancing personalized oncology and improving patient outcomes.

## Introduction

Therapy selectivity, defined as the ability of a drug to bind specifically to its intended target, is a critical aspect of the drug development pipeline. Many therapies fail in clinical trials due to side effects caused by off-target binding – or, in other words, lack of selectivity^1^. While the treatment of most diseases primarily requires target selectivity, other features may be crucial, such as tissue selectivity or, in the case of anti-cancer treatments, cancer-cell selectivity^2,3^. Target selectivity is related to the pharmacological properties of the drug, whereas tissue selectivity is associated with its pharmacokinetic properties^4^. Cancer-cell-selectivity involves specifically attacking targets predominantly present in cancer cells while sparing healthy cells^5^. In combination, the optimization of these selectivity features is crucial to enabling efficient and safe therapies^4,6^.

The advent of targeted anti-cancer therapies, also referred to as personalized or precision oncology, represents a significant shift in cancer treatment strategies and is based on the concept of therapy selectivity^7^. Compared to traditional cancer treatments such as chemotherapy and radiation therapy, targeted therapies leverage the unique genetic makeup of tumors to selectively target cancer cells^7^. Some of the characteristics of tumors that make this possible include the differential expression and the mutation of certain targets compared to their counterparts in healthy cells^8,9^. Protein kinases are predominantly used as personalized anti-cancer targets given their high relevance in cancer signaling and aberrant genetic landscape across cancer types^10,11^. In particular, mutated kinases may present functional or structural differences that distinguish them from their otherwise highly conserved protein family and that can be selectively targeted^12,13^.

Current anti-cancer kinase targets commonly overexpressed in tumor tissue include human epidermal growth factor receptor 2 (HER2) in breast cancer and fms-like tyrosine kinase 3 (FLT3) in acute myeloid leukemia^8^. Mutated anti-cancer kinase targets leveraged in the clinic exhibit distinguishing features, such as the fusion protein BCR-ABL present in most patients with chronic myeloid leukemia and resulting from the aberrant coupling of the genes of breakpoint cluster region protein (BCR) and tyrosine-protein kinase ABL1^14^. Other structurally distinguishing features of mutated anti-cancer targets that promote selectivity include altered orthosteric binding pocket conformations and novel allosteric binding pocket formation^15,16^. These have also been leveraged in the clinic, as exemplified by the epidermal growth factor receptor 1 (EGFR) L858R activating mutation targeted by selective orthosteric small molecule tyrosine kinase inhibitors (TKI)^17^ and clinical candidates targeting an allosteric pocket selectively in phosphoinositide 3-kinase α (PI3Kα) activating mutants^18^.

To make personalized oncology more accessible, the scientific community is actively engaged in expanding the range of druggable (mutated) targets^19^. However, several challenges arise regarding the criteria for identifying suitable candidates^20^. As previously discussed, the target should exhibit distinct characteristics compared to its healthy tissue counterpart to improve selectivity. Additionally, the candidate must play a functionally significant role in cancer progression to enhance efficacy. Lastly, the target should be relevant to a large group of patients with the cancer subtype. To achieve this, it is necessary to conduct various experiments that delve into potential candidates, covering structural, functional, and multi-omics analyses – a process that can be quite time- and cost-intensive^21^. In this context, different holistic computational strategies have emerged as particularly effective in compiling heterogeneous data types and prioritizing target candidates^22–25^.

Knowledge graphs, or complex networks, stand out as significant computational tools employed in cancer research due to their high versatility and interpretability^26^. By adopting graph-based data representations, these methods facilitate the storage and comprehensive analysis of diverse data entities (nodes) and their interrelations (edges). This functionality not only enables explicit knowledge retrieval but also facilitates the exploration and prediction of implicit knowledge by applying complex network algorithm analyses or deep learning approaches^27^. Some of the most fundamental graph-data representations in cancer research are gene regulatory networks and protein-protein interaction (PPI) networks, which represent causal or physical associations between entities of the same data type^28^. The nodes in these networks can be enriched with other types of data, such as clinical-relevant genomic data^29^, transcriptomics^30–32^, multi-omics^33^, disease-associated scores^32^, or inhibitor profiling^32^, and also combined with other networks^34,35^. Alternatively, knowledge graphs can be constructed with heterogeneous data nodes representing entities other than genes or proteins, such as genetic variants^36^, diseases^36–39^, phenotypes^38,40,41^, symptoms^39^, treatments^41^, drugs^36,37,40,41^, and risk and prevention factors^39^. These graphs have a broad range of applications, spanning from cancer diagnosis and subtype classification^35^ to prevention^41^ and treatment planning^40^. Among these applications there are also various tasks related to oncological drug discovery such as pathogenesis analysis^38^, mutant driver^34^ and resistance^29^ prediction, biomarker^30^ and target^33^ identification, drug repurposing^37^, and drug sensitivity prediction^23,31^. However, the multi-faceted nature of anti-cancer drug selectivity, which makes it an intriguing subject for study as a knowledge graph, remains largely unexplored in this context.

In this study, we introduce a knowledge graph approach for prioritizing cancer mutations with clinical, functional, and structural significance as potential targets for selective anti-cancer therapies. Due to limitations in data availability, our focus was on the human kinome, representing the complete set of protein kinases encoded in the human genome. This knowledge graph was constructed by integrating two distinct networks. Firstly, a pre-existing PPI network was used, linking protein-encoding genes through phosphorylation events, which are the main kinase signaling events^42^. Secondly, a patient-centric network was developed, connecting kinase somatic mutations based on their co-occurrence in cancer patients sourced from the Genomic Data Commons (GDC) database^43^. To enhance clinical and functional relevance, gene nodes in the knowledge graph were annotated with transcriptomics data.

Additionally, mutation nodes underwent structural and functional annotation through analyses from primary sources including the protein data bank (PDB)^44^ and KLIFS^45^. Finally, all nodes were enriched with bioactivity data from ChEMBL to evaluate druggability and drug sensitivity of mutations^46^. This comprehensive network serves as a valuable resource in cancer research, potentially facilitating the identification of relevant mutations for targeted therapeutic interventions.

## Results and discussion

### Overview of primary sources: clinical, structural, and functional relevance

A knowledge graph was constructed to enable the prioritization of kinase cancer mutations as selective targets for anti-cancer therapies. The perfect candidate was defined as a mutation of clinical relevance with the maximum potential to induce differential pharmacological effects compared to its wild-type counterpart and negligible potential to develop resistance mechanisms^47^. Clinical relevance was defined by a mutation recurrence in multiple cancer patients, but also by the kinase overexpression in a particular cancer type, which is a common druggability and cancer-cell selectivity indicator in targeted anti-cancer therapies. The last two conditions were further linked to the mutation’s structural and functional relevance. In this context, orthosteric ligand binding pocket mutations were considered structurally relevant due to their potential to modify the pocket’s conformation, which can be exploited to promote selectivity. These mutations were also considered functionally relevant because targeting them has the potential to directly disrupt the protein’s function. Moreover, additional resistance mutations in the pocket have the risk of disrupting binding of endogenous substrates needed for their activation and have thus a higher resistance threshold. Allosteric pockets have also been targeted in the past to increase target selectivity in cancer cells^15^, but they were not directly considered here due to constraints locating them. Functional relevance was also characterized by the importance of the target kinase in the cellular phosphorylation network. Apart from being central to crosstalk in cancer, kinases with a central role are less likely to have their signaling network re-routed and are therefore less prone to developing resistance^48^.

Data was therefore collected from primary sources across five layers of information to address the key questions guiding the selection of selective mutation target candidates with clinical, structural, and functional relevance (**Figure 1**). The information pertained to either mutations (top three layers in **Figure 1**) or their corresponding targets (bottom two layers in **Figure 1**). Kinase mutations were derived from cancer somatic mutation data from the NIH GDC dataset^43,49^ and their connections enabled the analysis of mutation co-occurrence in a patient and overall mutation recurrence (third layer). Mutations were further annotated with information regarding their location on the protein (second layer) and their pharmacological effect with respect to the wild-type protein (first layer). Structural location was determined through the analysis of data from a family-independent source (PDB)^44^ and a kinome-specific source (KLIFS)^45^. The analysis of PDB complexes enabled the annotation of the distance between the mutated residue and the ligand centroid as a proxy for the distance to the orthosteric binding site, as previously described^50^. Additionally, the aligned position in the kinase binding pocket was annotated from KLIFS. The pharmacological effect of mutations was annotated from the combined analysis of the differences between bioactivity distributions in mutant and wild-type targets in ChEMBL and the Papyrus dataset, as previously described^50^. Apart from the kinases with cancer-related mutations, targets were derived from the phosphorylation PPI network defined by Olow *et al*.^42^ and their connections supported the analysis of phosphorylation events between kinases and their substrates (fourth layer). Genes encoding the protein targets were further annotated with their differential expression in different cancer types compared to normal tissue that was derived from the GDC dataset (fifth layer).

**Figure 1.**
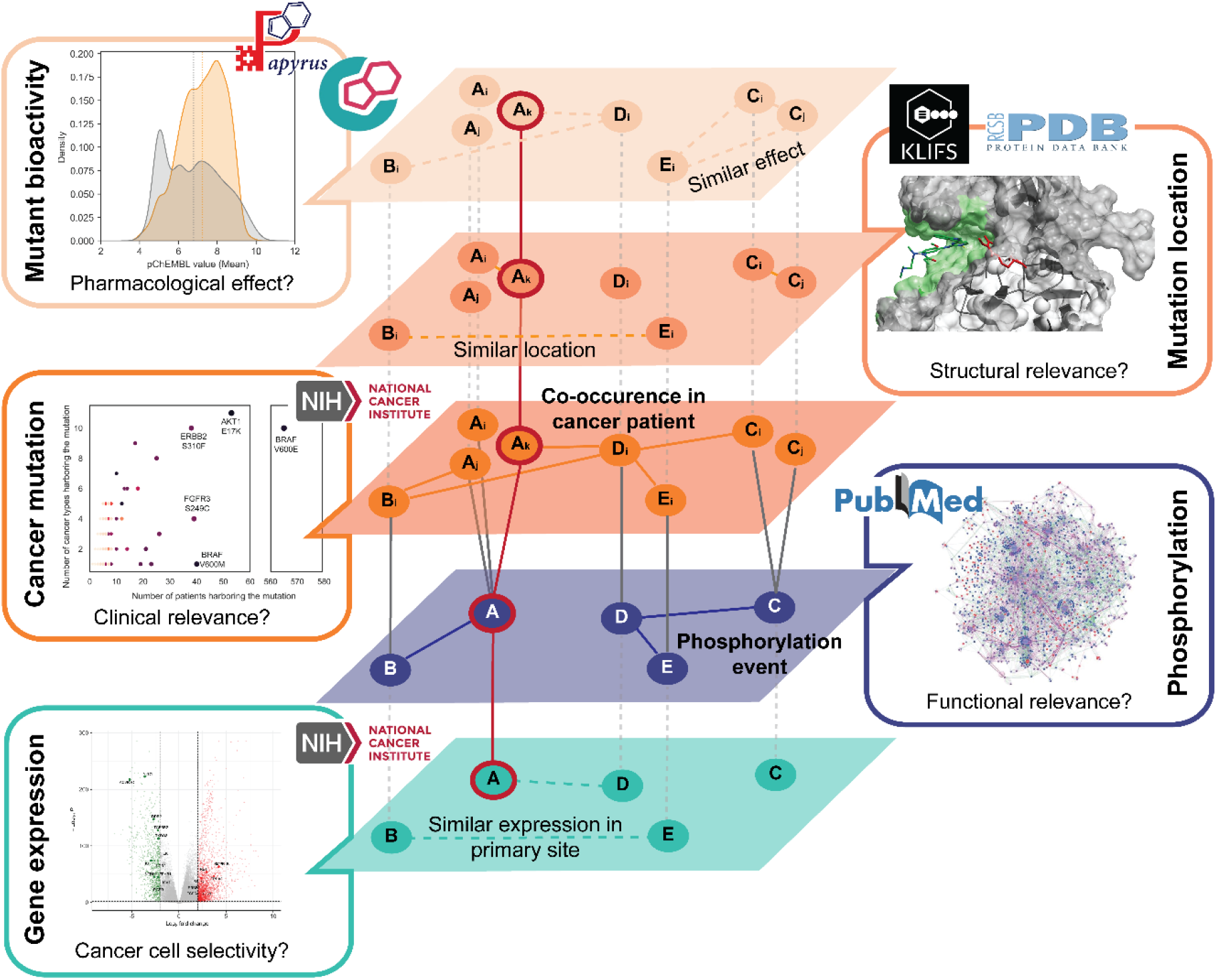
Overview of five layers of information contained in the knowledge graph. The top three layers correspond to somatic mutations, while the two bottom layers correspond to gene / protein targets. From top to bottom, the first layer represents the bioactivity differences triggered by mutations with respect to wild-type. The second layer represents the mutation’s structural location in the target protein. The third layer represents the occurrence of mutations in cancer patients. The fourth layer represents the phosphorylation events between targets. Finally, the fifth layer represents the differential expression of the target genes tumor tissue in different cancer types (defined by the primary site where the tumor developed) compared to normal tissue. Each layer answers a distinct question about the mutation’s relevance as an anticancer target. The connections represented by lines between entities (nodes) within the third and fourth layers are used as edges in the knowledge graph, as well as the connections between nodes of those two layers. The connections within the rest of the layers, represented by dashed lines, are not kept in the knowledge graph but indicate how nodes relate based on the data represented in each layer. The red line connecting and highlighting nodes across the five layers exemplifies a potential candidate with information spawning across the five layers.

While knowledge graphs can incorporate a larger variety of information, such as Astra Zeneca’s BIKG that was constructed with 37 public and internal datasets^51^, subsets of the graphs^29,52^ or more tailored representations^32^ are needed to answer specific research questions. For example, the CancerOmicsNet graph was created with the purpose of predicting therapeutic effects in various cancer cell lines. It incorporated layers of information similar to those mentioned here, consolidated into a single graph using a phosphorylation PPI network^32^. In contrast, our approach involves integrating mutation-specific data that aligns with the specific objectives set for this graph. The data across these five layers enables an investigation into the clinical, structural, and functional implications of mutations and targets, while also tackling important considerations around druggability, like selectivity towards cancer cells and mutations. This information was incorporated into a knowledge graph and analyzed accordingly, as detailed in the following sections.

### Knowledge graph architecture

Mutations and targets were connected in the knowledge graph by association edges and comprised the two types of nodes available. Mutation nodes were connected by edges representing co-occurrence in the same cancer patient. Target nodes – referred to as gene nodes from now on to denotate target protein-coding genes - were connected by edges representing all phosphorylation events between kinases and their substrates (**Figure 2a**). The rest of the information collected from primary sources was collapsed from the five layers of information into one and stored as – mutation or gene – node attributes (**Figure 2b**). Edges representing cancer patient co-occurrence were also annotated with attributes representing the corresponding patient and cancer type, which allowed analysis per cancer type in subsequent sections. This simple graph architecture was chosen to keep mutations and their corresponding protein coding genes as the central elements. The knowledge graph was constructed in NetworkX^53^ allowing multiple connections between the same two nodes. In the final kinome graph this amounted to 78,782 nodes and 6,515,059 edges. Node entities and their relationships were determined manually here to maximize accuracy. However, other options are becoming available to extract them automatically, such as using large language models^54^, in turn facilitating ontology mapping^39^. These novel approaches are particularly relevant and promising in large knowledge graphs comprising numerous primary sources, although several challenges still need to be addressed, particularly regarding precision and biases in the data extracted^55^.

**Figure 2.**
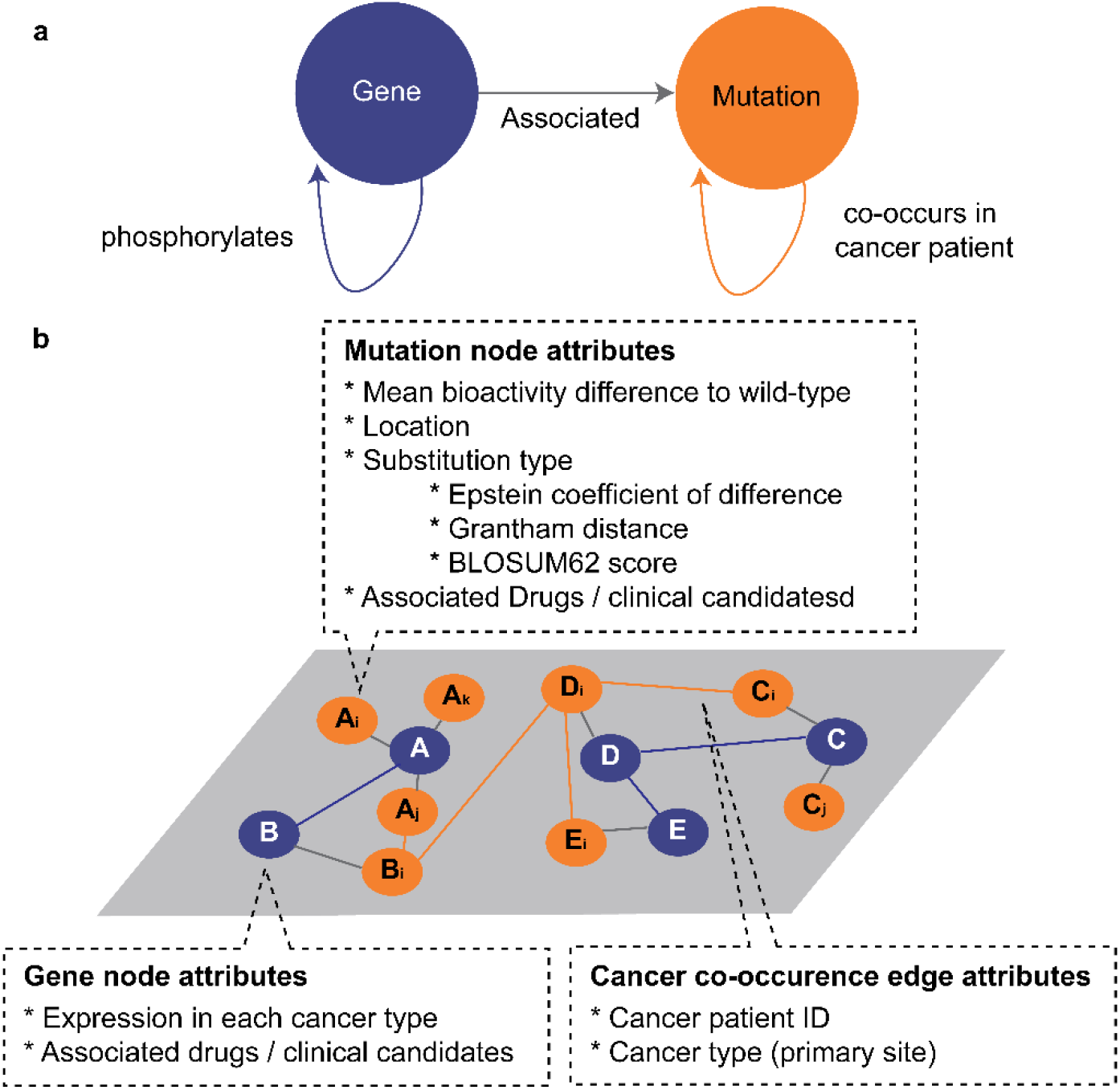
Knowledge graph architecture. **a)** Schematic representation of the two types of nodes in the graph (circles) and their edges (arrows). **b)** Example graph representation with attributes linked to mutation and gene nodes, as well as to cancer co-occurrence edges. Mutation nodes are represented in orange and gene nodes in blue. Edges connecting different types of nodes correspond to the edges described in **a)**.

The distribution of types and subtypes of nodes and edges is presented in **Table 1**. Gene nodes amounted to 1,571, of which 667 were kinases and 904 substrates. Of note, the original phosphorylation PPI network contained 774 kinase nodes but only 625 were kept after applying the filter for proteins with kinase activity as defined in the methods section. Additionally, 42 genes coding for proteins with kinase activity that were not included in the original phosphorylation PPI network were added to the graph due to the presence of mutations in cancer patients. Substrate types were further investigated, particularly within the context of membrane proteins. Specifically, 17 nodes were identified as G protein-coupled receptors (GPCRs), while 14 were categorized as solute carriers (SLCs). Additionally, 130 other substrates were found to be primarily localized to the plasma membrane. Moreover, out of the kinase nodes, 65 were also receptors. The majority of nodes (77,009) represented cancer mutations. Out of these, only 34 were associated with bioactivity data from both ChEMBL and the Papyrus dataset. Consequently, an additional 202 kinase mutation nodes with bioactivity data were incorporated into the graph to allow the interpolation of mutation characteristics affecting bioactivity. In an expanded version of the graph aimed at drug discovery other node types could include small molecules^36,37,40^, instead of summarizing the effects of mutations over all tested drugs. This could result in an additional layer of information where edges represent similarity between small molecules^56^. An alternative would be to report the effect of individual kinase inhibitors as individual attributes in each kinase node^32^. However, other primary sources for mutation-driven bioactivity changes should be considered before implementing any of these expansions^57^.

**Table 1.** Distribution of node and edge types and subtypes across the kinome knowledge graph.

| Entity | Type | Subtype | Number of entities |
| --- | --- | --- | --- |
| Nodes | Gene | Kinase | 667 |
|  |  | Substrate | 904 |
|  | Mutation | Cancer | 77,009 |
|  |  | Other (ChEMBL + Papyrus) | 202 |
| Edges | Gene - Gene | Phosphorylation | 5,759 |
|  | Gene - Mutation | Cancer | 87,248 |
|  |  | Other (ChEMBL + Papyrus) | 202 |
|  | Mutation - Mutation | Cancer patient co-occurrence | 6,421,798 |
|  |  | Other (ChEMBL + Papyrus multiple substitutions) | 52 |

Filtering nodes from the original phosphorylation network resulted in a reduction of phosphorylation edges between gene nodes from the original count of 5,963 to 5,759. Cancer mutation nodes were connected to their respective genes via one or multiple edges, indicating the frequency of the mutation within that gene across patients. Cancer mutations were collected from 8,518 patients across 48 cancer types. As a result, the number of edges between cancer mutations and genes exceeded the count of cancer mutation nodes by 10,239. Conversely, the number of edges connecting genes and non-cancer mutations equaled the count of non-cancer mutation nodes. Over 6.4 million edges represented the co-occurrence of two mutations in the same cancer patient, highlighting a high mutation burden in kinases. The distribution varied widely, with some patients having just one edge representing one mutation and others having up to 672,220 edges representing up to 1,159 unique mutations. The median number of co-occurring edges per patient was six, indicating most patients had few kinase mutations, while a few had a large number of mutations.

The analysis of node degrees in the knowledge graph confirmed this irregular pattern, highlighting the presence of a few nodes with exceptionally high degrees and a much larger number of nodes with lower degrees (**Figure 3**). A node degree represents the number of edges linking it to other nodes. As anticipated, recurrent cancer mutations and their corresponding genes exhibit high degrees within the knowledge graph. PIK3CA R88Q and BRAF V600E are the two mutations with the highest degree (10,275 and 6,219, respectively) and were present in 68 and 565 patients respectively (**Supplementary Table 1, 2**). Interestingly, other PIK3CA mutations with a higher occurrence frequency in cancer patients (PIK3CA E545K and H1047R present in 258 and 234 patients, respectively) showed comparatively lower node degrees (2,484 and 2,439, respectively). These results emphasize the importance of mutation co-occurrence and tumor mutation burden (TMB), which has recently been highlighted as a tumor biomarker^58^. Accordingly, genes linked to TMB, such as TTN and OBSCN^59^, were also highlighted in the knowledge graph based on their node degree. In a similar manner, mutations not highly recurrent across cancer patients but present in patients with a high TMB across cancer types (**Supplementary Table 3**) showed a high node degree, three of them even being present in natural variance (**Supplementary Table 2**). Mutation co-occurrence in patients has also been highlighted in other graph approaches independent of mutation recurrence^60^. However, in order to pinpoint exclusively to functionally relevant mutations, it might be appropriate to filter frequent mutations *a priori*^61^. From a graph architectural point of view, the emphasis on individual patients that is central to our approach has also been reported in other patient-centric graphs. The most common approaches also consider genes in PPI networks that are enriched with multi-omics data^62,63^. However, other graph architectures are possible where nodes represent individual patients with different multi-omics attributes^64^. Incorporating patient nodes into the existing graph and linking them to their corresponding mutations and genes could aid in filtering out patients with high TMB and mutations with low recurrence rates, resulting in a more refined graph. As elaborated in the subsequent sections, addressing data sparsity within the graph is a significant bottleneck for analysis, and constructing focused subgraphs can sometimes mitigate this issue.

**Figure 3.**
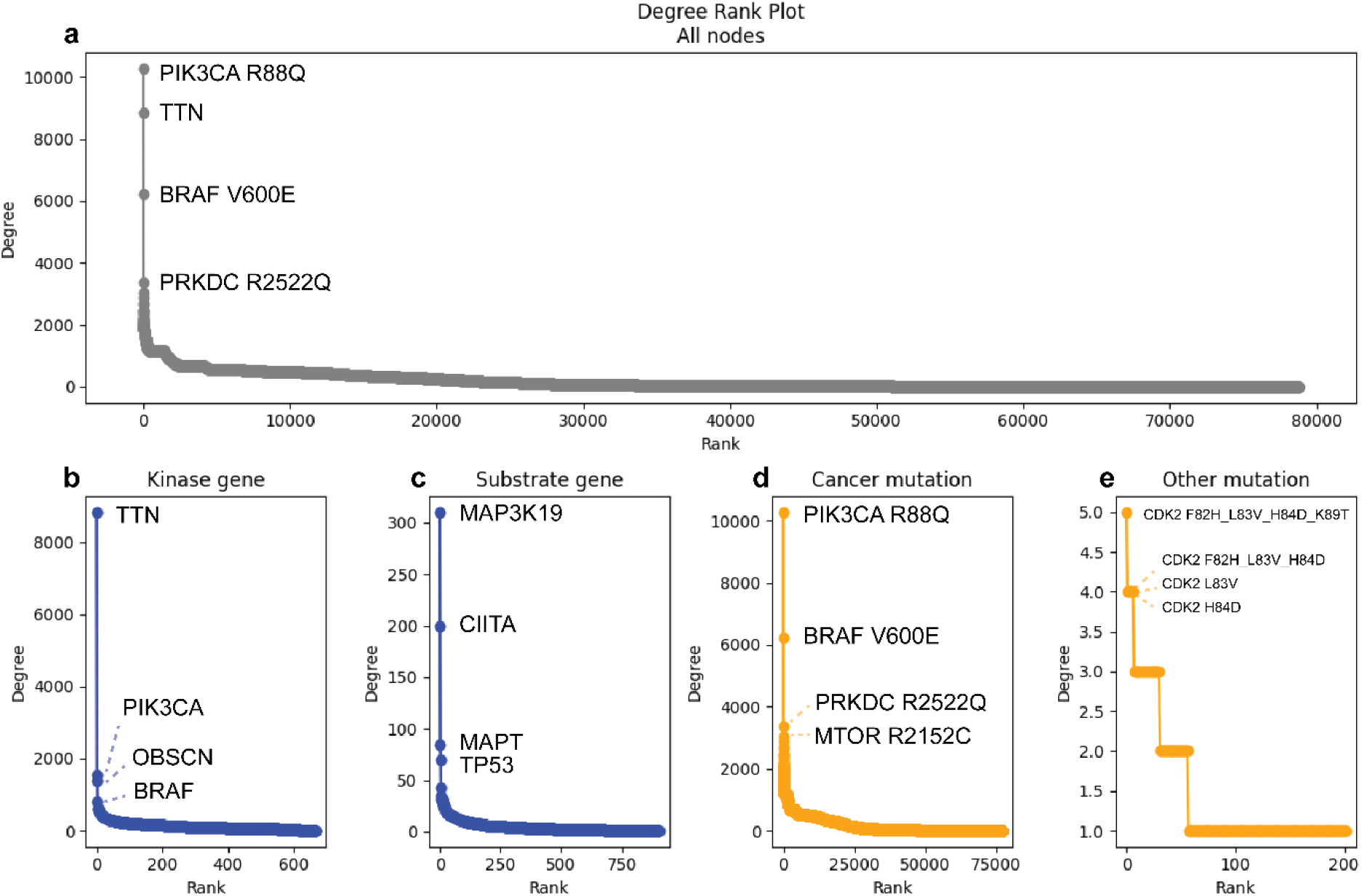
Node degree rank analysis in the kinome knowledge graph. Nodes are ranked based on their degree, which is calculated as the number of edges connecting the node to other nodes in the graph. The degree rank analysis is calculated for all nodes in the graph (**a**) as well as for each node subtype independently: kinase (**b**) and substrate (**c**) gene nodes, and cancer (**d**) and other (**e**) mutations. The top four ranked nodes in each case are labeled accordingly.

### Attribute annotation sparsity across the graph

Nodes in the graph representing mutations and genes were annotated with attributes covering the information across the five layers of information previously described in **Figure 1**. Node attributes are crucial for graph analysis, as they can further denote the embeddings or labels for predictions in machine learning applications. In particular, the number of approved drugs and bioactivity changes were key mutation attributes that could be used to classify mutations for targeted therapy. However, the annotation density (this is, how many nodes have non-null values for a particular attribute) of these two and many other node attributes was rather low (as depicted in **Figure 4** for mutation nodes, and in **Supplementary Figure 1** for gene nodes).

**Figure 4.**
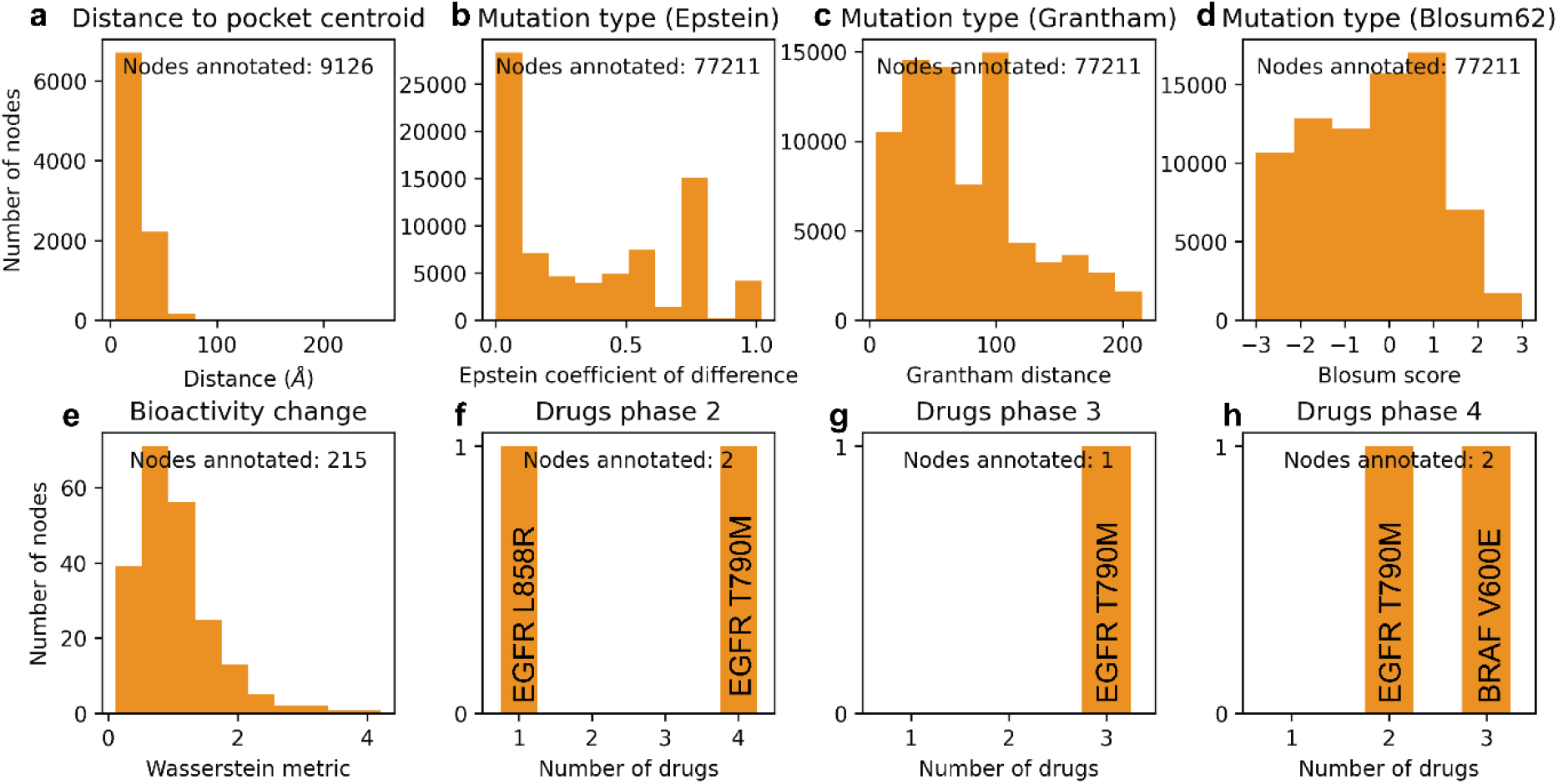
Density and distribution of mutation node attributes in the kinome knowledge graph. For each of the attributes, the number of mutation nodes with non-null values for the particular attribute is depicted on the y-axis. Moreover, the graphs represent the distribution of the attribute values across the mutation nodes on the x-axis, in the form of histograms or bar plots, depending on the density of each attribute. Distribution is represented as histograms for the distance to the pocket centroid calculated from PDB complexes (**a**), the mutation type as determined by the Epstein coefficient of difference (**b**), the mutation type as described by the Grantham distance (**c**), the evolutionary probability of the mutation type as described by the Blosum score in the Blosum62 matrix (**d**), and the bioactivity change represented by the Wasserstein distance between the bioactivity distribution for the mutation and the wild-type protein found in ChEMBL (**e**). Bar plots represent the number of drugs in different phases of development according to ChEMBL labeling: clinical phases 2-3 (**f**-**g**), and approved drugs (**h**). Pre-clinical candidates (phase 0) and drugs in clinical phase 1 are not included because they were not annotated in any mutation nodes. The mutations represented in the bar plots are labeled for reference.

Of the 1,571 gene nodes in the graph, 351 (22.34%) have at least one drug in any phase of development directly linked on ChEMBL as its target (**Supplementary Figure 1**). However, only three kinase mutations are listed as the target for drugs approved or in development (**Figure 4f-h**). These are BRAF V600E and EGFR L858R activating mutations and EGFR T790M acquired resistance mutation, all previously linked to cancer. Apart from being too low to generate labels for classification tasks, these annotations are a direct consequence of the non-triviality of the variant annotation pipeline in bioactivity databases that was previously described^50^. For example, EGFR L858R does not have any approved drug directly linked to the mutation in the mechanism of action, probably due to the fact that most approved first generation EGFR inhibitors were developed to target selectively either the activating deletion in exon 19 or the activating mutation L858R, and deletions are not fully curated in ChEMBL as of yet – although this is work in progress. Similarly, only 215 (0.28%) mutation nodes have a bioactivity change annotation (**Figure 4e**), which is a very small number for further machine learning analysis. These numbers could be increased with data from additional primary sources for mutant-induced bioactivity changes, such as the mutation-induced drug resistance database (MdrDB)^57^.

Mutation attributes representing the characteristics of the amino acid substitution that could be used for the node embeddings were less sparsely represented (**Figure 4a-d**). Mapping of the three metrics representing mutation types (Epstein coefficient of difference^65^, Grantham distance^66^, and Blosum62 score^67^) could be done for all 100% of the mutations. These three metrics were selected to cover different aspects defining the amino acid substitutions, namely physicochemical properties (Epstein coefficient of difference, which is directional and represents the size and polarity difference between the wild-type and mutated amino acid; and Grantham distance, which is non-directional and calculates said difference based on atomic composition and molecular volume on top of polarity), and evolutionary conservation (Blosum score, which defines the evolutionary probability of a substitution relative to random probability, calculated for proteins clustered at 62% sequence similarity). Of note, 57% of the mutations had an Epstein coefficient of difference lower than 0.4, and 74% a Grantham distance below 100, meaning that the majority of substitutions were rather conservative. Simultaneously, 54% of the mutations were likely to happen by chance, as represented by a Blosum score of zero or higher, and in fact, 2,759 (3.57%) of the all the mutations were also found to occur as natural variance in the 1000 Genomes dataset previously compiled^24^. The complementarity of the three amino acid substitution metrics was illustrated by the variations observed in their distributions **(Figure 4b-d)** while still aligning with clinical significance (**Figure 5**). From the 19 most recurrent mutations in cancer patients (occurring in more than 20 patients), six mutations (BRAF V600E, FGFR3 S249C, ERBB2 S310F, FGFR2 S252W, EGFR L858R, and PIK3CA C420R) were identified as disruptive based on an Epstein coefficient of difference over 0.4 and a Grantham distance greater than 100. Additionally, these mutations were determined to be less likely to occur by random chance based on a negative Blosum score. The most common mutation, BRAF V600E, occurring in 565 patients, had an Epstein coefficient of difference of 1.0 (**Figure 5a**), a Grantham distance of 121 (**Figure 5b**), and a Blosum score of -2 (**Figure 5c**). It is key to note, however, that three oncogenic PIK3CA mutations highly recurrent in breast cancer (E545K, H1047R, and E542K^68^) would not be captured by any of these three metrics, while the three mutations with associated drugs under development (BRAF V600E, EGFR L858R, EGFR T790M) would. Therefore, although these annotations are useful to get a general understanding of the potential effect of an amino acid substitution, more advanced amino acid embeddings may be preferred for machine learning applications ^69^.

**Figure 5.**
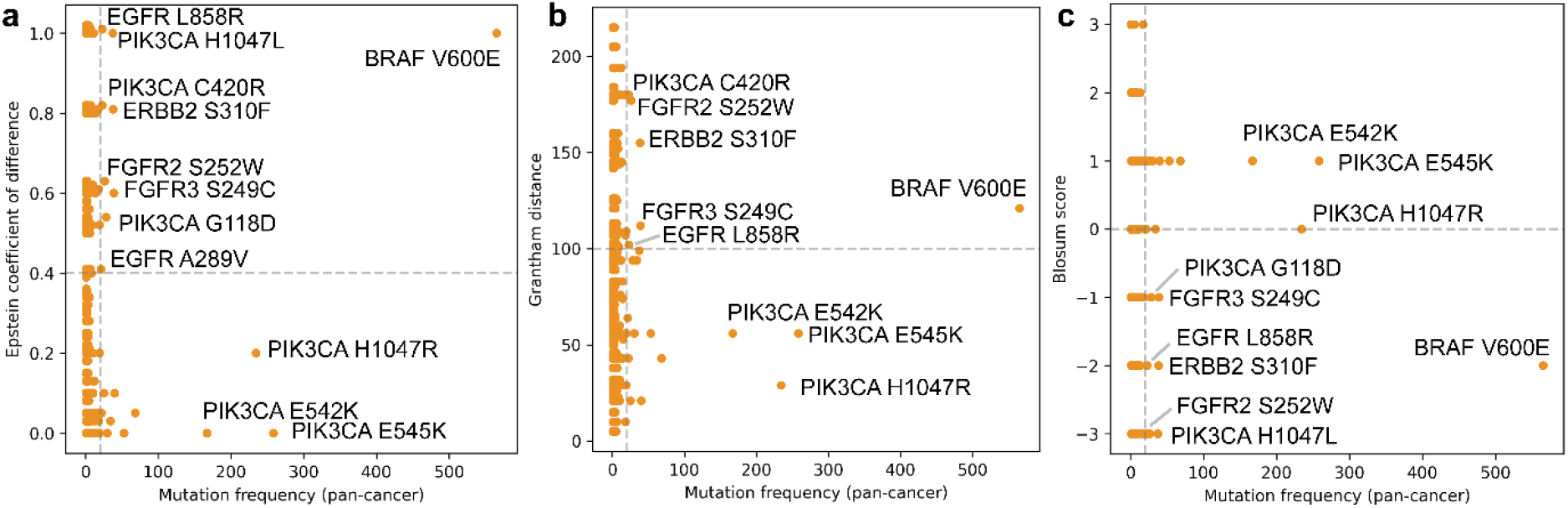
Correlation between cancer mutation frequency and three metrics describing the amino acid substitution in the kinome knowledge graph: Epstein coefficient of difference (**a**), Grantham distance (**b**), and Blosum score (**c**).. Mutations occurring in more than 100 patients pan-cancer are labeled for reference. In **a)**, an Epstein coefficient of difference of 0.4 is taken as an arbitrary threshold to distinguish between conservative (<0.4) and disruptive substitutions (>0.4), which are labeled for reference if they occur in more than 20 patients pan-cancer. In **b)**, a Grantham distance of 100 is taken as an arbitrary threshold to distinguish between conservative (<100) and disruptive substitutions (>100), which are labeled for reference if they occur in more than 20 patients pancancer. In **c)**, substitutions with an alignment happening less often than random chance as collected in the Blosum62 matrix (Blosum score < 0) are labeled for reference if they occur in more than 20 patients pan-cancer.

Attributes representing the structural location of the mutation in the protein were mapped to 9,126 mutations (11.82%) occurring in 161 genes when calculated from PDB complexes (**Figure 4**), and 3,854 mutations (4.99%) occurring in 295 genes were annotated with a KLIFS pocket position (**Supplementary Figure 2a**). Interestingly, only 1,890 mutations (2.44%) occurring in 138 kinases had a double structural annotation (from PDB and KLIFS), which highlighted the complementarity of these approaches. The annotations extracted from KLIFS covered a larger number of kinases, since they also included pocket annotations for kinases with only apo structures available in the PDB, such as TTN. An additional advantage of the KLIFS annotation is that it can be directly linked to its structural relevance – functionally and pharmacologically. The PDB annotations, on the other hand, enabled the annotation of mutations in an additional 22 genes for which there was no data available in KLIFS. More importantly, they enabled the annotation of mutations outside of the ATP binding pocket, providing a more holistic view of the structural location of the mutations, which is very relevant for drug response prediction^70^. In fact, while secondary resistance mutations tend to be part of the ATP binding site, many activating mutations crucial for cancer development are outside of the KLIFS-defined binding pocket. For example, from the 25 most frequent cancer mutations shown in **Supplementary Table 1** and **Figure 5**, only EGFR L858R is part of the KLIFS binding site. This is also exemplified by the discrepancy identified between the number of mutations within the KLIFS binding pocket and the maximum mutation frequency at those positions (**Supplementary Figure 2a,c**). While positions c.l.69 and c.l.74 in the kinase catalytic loop were the most frequently mutated overall with 91 and 88 individual mutations respectively, the two most frequent individual mutations in cancer patients within the pocket were EGFR L858R in the activation loop (a.l.84) and ERBB2 V842I in the β-sheet VI (VI.67), occurring in 23 and 17 patients, respectively. It is worth noting that none of the seven binding pocket positions with mutations occurring in 10 or more patients were highly conserved, as reported by KLIFS. This highlights the need for cancer cells to not fully disrupt kinase function^71^. The analysis of mutations annotated with both structural sources allowed for the correlation of the KLIFS pocket positions with precise distances to the ligand centroid. This demonstrated variability in the distances that is consistent with varying ligand sizes and binding modes observed among kinases (**Supplementary Figure 3**). However, this analysis also highlighted the presence of outliers representing measurement errors, which may arise from inconsistencies in sequence numbering and should be corrected in the pipeline. For kinases, one solution could be to utilize the KLIFS-curated structures, although this approach is specific to the kinome and cannot be expanded to other protein families. Other solutions are possible to structurally annotate cancer mutations and incorporate this information to PPI networks, but the existence of incomplete and incorrect structures in the PDB is a constant bottleneck^72^.

In general, the sparsity of knowledge graphs is a significant bottleneck for machine learning applications as it compromises the quality of the embedding methods used *a posteriori*^73^. However, sparsity in node attributes may not be a problem if the bias is in sync with the graph’s objective and can be utilized for additional analysis. For example, some applications use customized attention mechanisms to prioritize nodes with more information for efficient data propagation to their neighbours^23^. Similar approaches could be used in the knowledge graph developed here to predict bioactivity differences or approved drug development for mutations using node attribute competition tasks, where node attributes are inferred from the collective data in the graph^74^. While some of the methods used for node attribute competition tasks are still robust with high node attribute sparsity (up to 80%)^74,75^, a minimum might still be required. It is important to note that some knowledge graphs are not necessarily developed with the aim to apply additional (machine learning) analyses, but with the intention to be a resource that can be interactively explored to gain a holistic view of a certain problem^36,39^. In the last section, we explore the development and analysis of different subsets of the kinome knowledge graph, which can be a strategy both to decrease node attribute sparsity with the aim to apply machine learning analyses, as well as to create biologically relevant subgraphs that can be interactively explored for data extraction.

### Subgraph analysis and interactive exploration: A case study for RTKs

The architecture of the knowledge graph supported the construction of biologically relevant subgraphs based on node and edge attributes. Although the python package developed to build and analyze the graph supports filtering the graph for any attribute of interest, two main types of subgraphs were pre-defined and explored in this section. To illustrate these analysis options, a smaller graph was created specifically focusing on receptor tyrosine kinases (RTKs) and their substrates, mimicking the structure of the kinome graph (**Supplementary Tables 4-6** and **Supplementary Figures 4-8**). The smaller graph was built to facilitate analysis and interactive visualization, but it also served as a way to increase node attribute density while maintaining biological relevance. Compared to the kinome graph, the RTK graph contained approximately a seventh of the original nodes (11,989 nodes instead of 77,211) and almost 50 times fewer edges (141,311 instead of 6,515,059). All mutation node attributes considered in the previous section remained sparse, but in all cases the percentage of mutation nodes annotated increased. For example, the percentage of mutation nodes annotated with bioactivity data increased from 0.28% to 0.77%, and the number of nodes annotated with distance data increased from 11.82% to 15.31% for PDB-annotated nodes and from 4.99% to 7.17% for KLIFS-annotated nodes. While modest, this rise in attribute density is consistent with the significant emphasis on cancer research related to RTKs^76^. The interactive visualization module enabled, among other views, the visualization of the phosphorylation edges in the RTK graph between the selected kinases and their substrates (dark and light blue, respectively in **Figure 6a**, where node size is proportional to the total number of drugs under development or approved for that node). To improve visualization, different node and edge types and subtypes can be left in the background, as was done in **Figure 6a** for mutation nodes and their edges. Of note, the RTK graph still contains some kinases that are not receptors because they are substrates of RTKs, such as JAK2 that is phosphorylated by EGFR. This can be easily explored by zooming into a particular node, as shown in **Figure 6b** for EGFR and all nodes connected to it. The zoomed-in view also enables the interactive exploration of node attributes, on the right-side panel, which for a gene node includes among others expression values for different cancer types.

**Figure 6.**
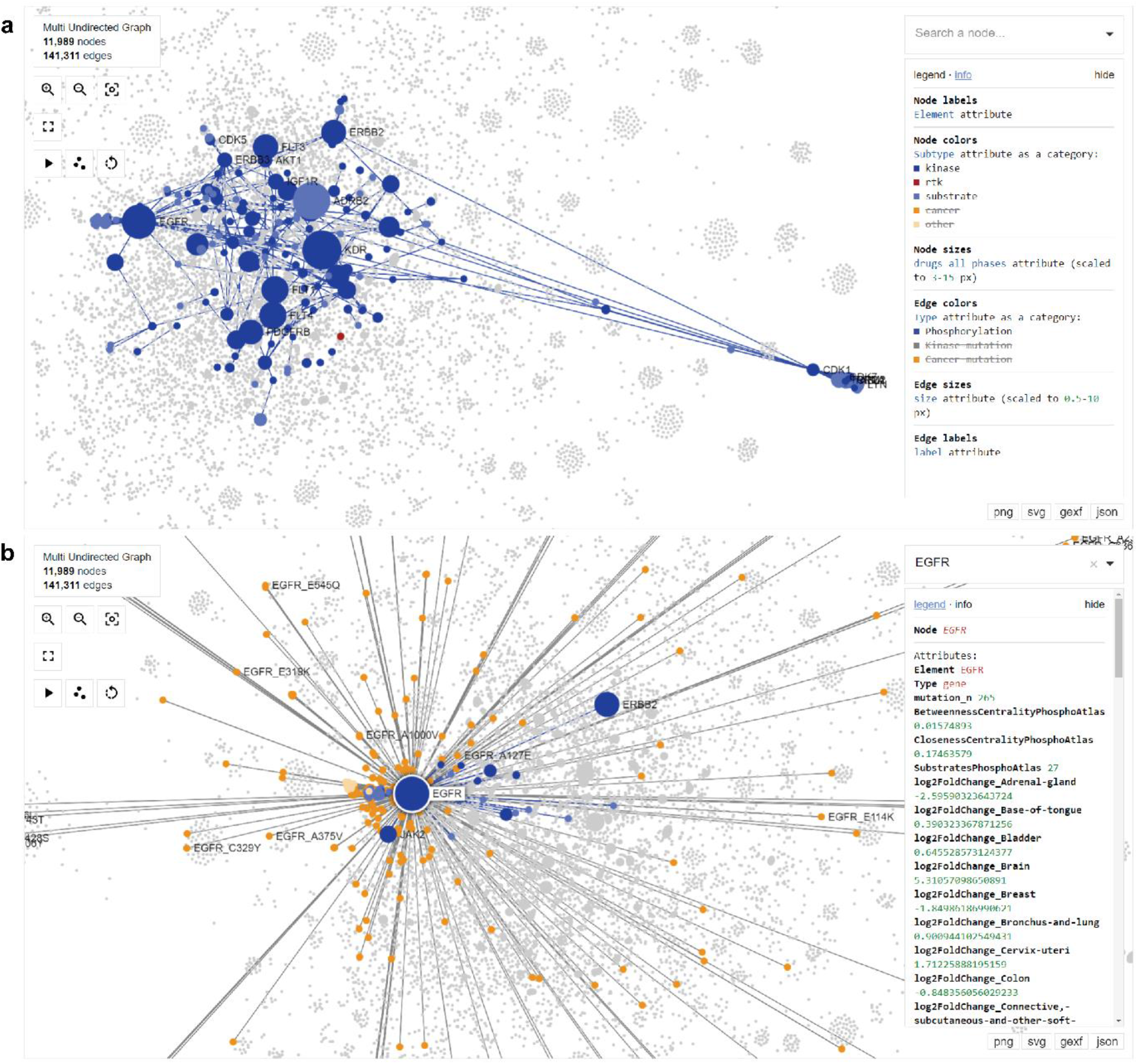
Interactive visualization example for the receptor tyrosine kinase knowledge graph. **a)** Visualization of the phosphorylation events between kinases (dark blue nodes) and their substrates (light blue nodes). Every highlighted edge represents a phosphorylation event. The visualization module, powered by IPysigma^77^, includes a legend panel on the right side that enables search options and filtering and describes the attributes used for node color and size. To improve visualization, all mutation nodes and edges are hidden here. **b)** Example visualization when clicking on a particular node, in this case, EGFR. All edges connecting the node of interest and the paired nodes are highlighted, while the rest of the graph is kept in the background. When a node is selected, all the attributes associated with it are displayed on the right-side panel.

The first of the two pre-defined analysis options supports the construction and exploration of individual “layer” subgraphs that are based on the type of edges. This enables the division of the knowledge graph into three subgraphs: a purely cancer patient-centric network linked by mutation co-occurrence, an association network between kinases and their mutations, and a phosphorylation PPI network. This division is important because it allows for the calculation of graph metrics for each layer independently. This, in turn, enables the characterization of the importance of specific nodes at various levels. By analyzing EGFR across multiple layers, it was possible to determine which layers influenced the final degree metric in the graph (**Figure 7a**). In this particular case, a high degree in gene nodes mostly arose from the kinase-mutation layer (**Figure 7b**) since there was an additional edge for each patient carrying a specific mutation. However, this high node degree could also result from a high degree in the phosphorylation layer (**Figure 7d**). High degree in mutation nodes, however, mostly resulted from mutation co-occurrence in the same patient (**Figure 7c**). While the results were consistent with the anticipated graph architecture, conducting an analysis of each layer reveals that EGFR exhibits a high degree of connectivity not only due to its prevalence in mutations across various patients, but also because of its multiple interaction partners within the phosphorylation network. This analysis can also be extended to other metrics, for example betweenness centrality (**Supplementary Figure 9**), which represents the influence a node has on the flow of information in the graph. This metric can in turn help determine different community clusters within the graph, as well as key nodes and edges relevant to connections between these communities.

**Figure 7.**
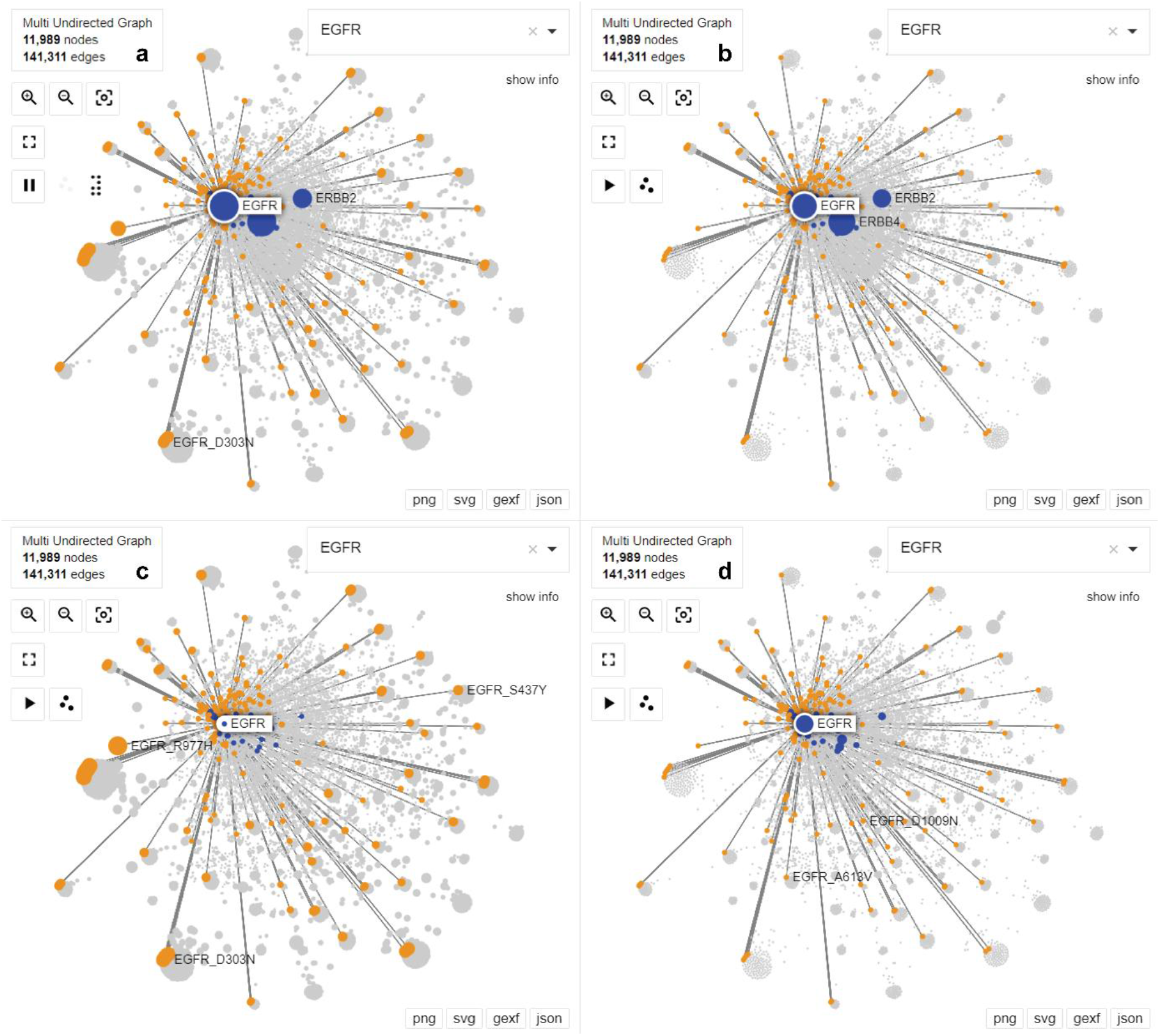
Node degree comparison across layers in the receptor tyrosine kinase knowledge graph, with focus on nodes connected to EGFR for visualization purposes, as done in Figure 6b. Gene nodes are represented in blue and mutation nodes are represented in orange. Nodes that are not connected to EGFR are kept in the background and represented in grey. Node size in each panel is determined by the node degree calculated from the whole graph (**a**) or one of the three pre-defined analysis layers: kinase-mutation layer (**b**), cancer-mutation co-occurrence layer (**c**), or phosphorylation layer (**d**).

The second pre-defined analysis module allowed individual subgraphs to be easily constructed for each cancer type by filtering the cancer-related edges corresponding to patients with a particular cancer type. These subgraphs also included the phosphorylation events pertinent to cancer-type filtered nodes. The RTK graph covered 44 cancer types, of which the six most populated cancer types were bronchus and lung (723 patients), skin (356 patients), brain (312 patients), corpus uteri (302 patients), bladder (289 patients), and breast (273 patients). The analysis of cancer type subgraphs enabled the distinction between recurrent and patient-specific mutations, since the former aggregate in the central part of the graph while the latter form clusters in the graph periphery. This phenomenon was clearly distinctive between the bronchus and lung subgraph (**Figure 8a**) and the skin subgraph (**Figure 8b**), where edges were colored to represent different patients. Most of the RTK mutations occurring in lung cancer patients were recurrent across patients, while several skin cancer patients harbored mutations that are specific for that patient. Similarly to the full graph, different node attributes can be used to interactively explore the graph. For example, the use of cancer type-specific gene expression (Log2 fold change) illustrated in **Figure 8** can aid in distinguishing genes with a higher relevance in that particular biological context. In fact, the construction of individual graphs for different genetic makeups is a common strategy when trying to prioritize cancer-related mutations^78,79^. Additionally, each subgraph can be analyzed independently and the results can be combined and displayed in the main graph, as done with the layer analysis. This further enables the investigation of cancer type-specific relationships in a pan-cancer context (**Supplementary Figure 10** for degree analysis and **Supplementary Figure 11** for differential expression analysis), which tends to be the preferred strategy in personalized anticancer therapies^80^.

**Figure 8.**
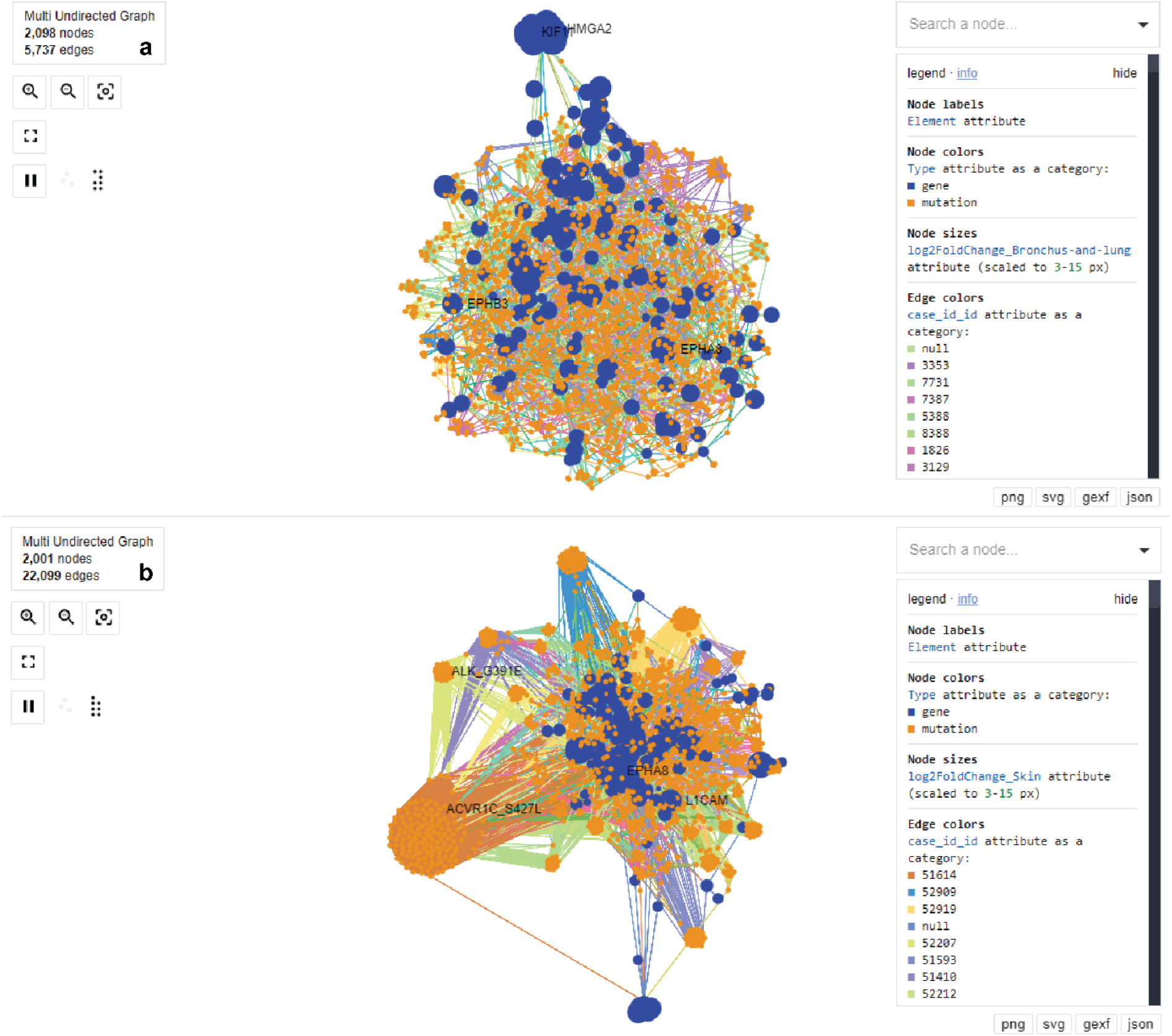
Interactive visualization of cancer type subgraphs derived from the receptor tyrosine kinase knowledge graph for the two most populated cancer types: bronchus and lung (**a**, 723 patients) and skin (**b**, 356 patients).Gene nodes are represented in blue and mutation nodes are represented in orange. Node size represents the differential expression (Log2 fold change) of genes in that cancer type tumor tissue compared to normal tissue. Each edge color represents a different patient.

**Figure 9.**
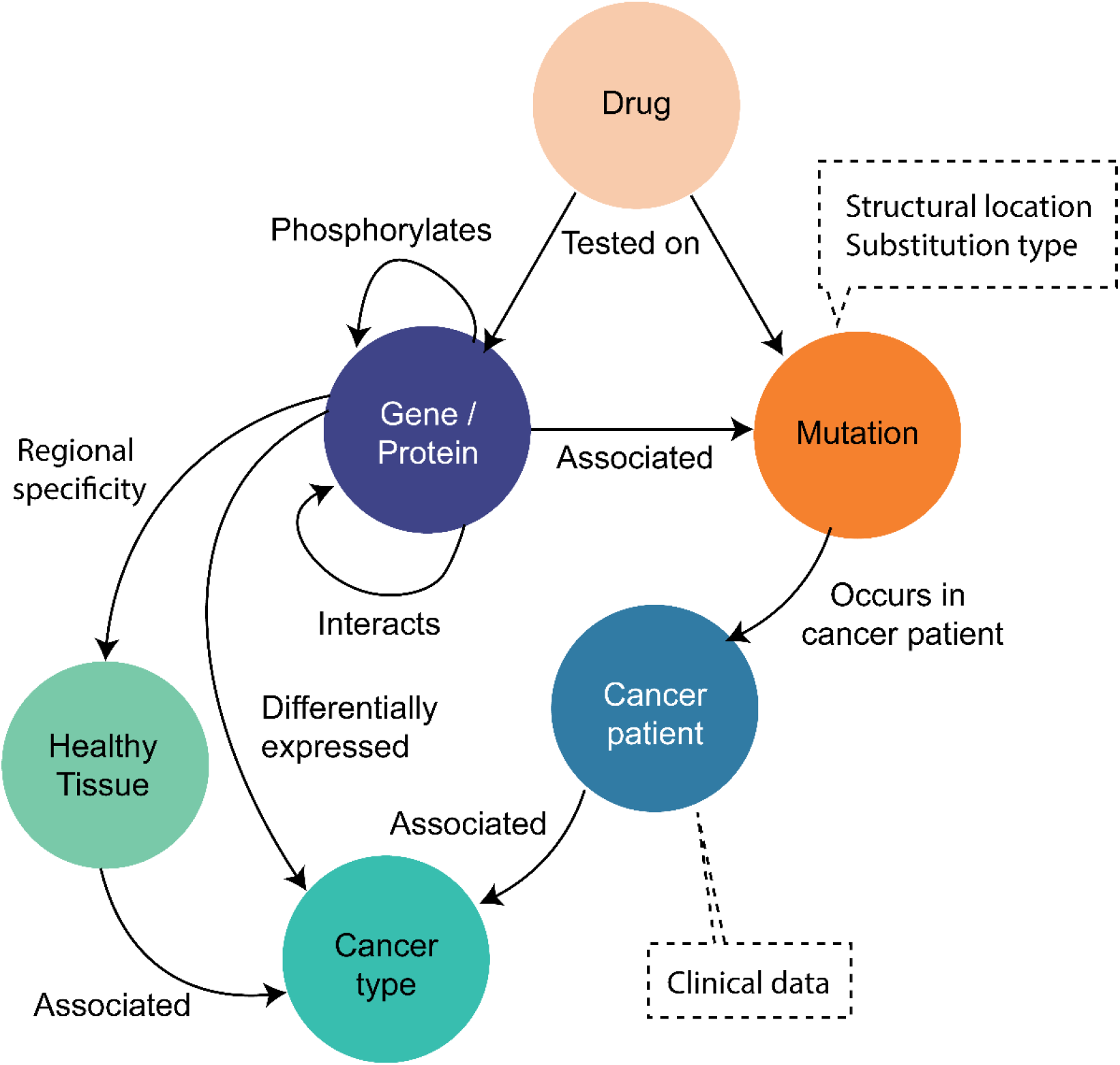
Proposed alternative knowledge graph architecture for the kinome patient-centric knowledge graph. Each circle represents a different node type and the arrows represent the edges between nodes. Dashed boxes include attributes that could be associated with specific types of nodes to increase information density.

## Conclusions

A cancer patient-centric knowledge graph was constructed to aid in identifying mutations with advantageous characteristics to be targeted selectively by small molecules, hence reducing side effects of anticancer therapies while maintaining high efficacy. Known oncogenic and targeted genes and mutations stand out from this exercise based on several of the node attributes collected across five layers of information, highlighting the potential of this graph in oncology drug discovery. Although initially tailored for the kinome, this framework can readily be adapted to other protein families by modifying kinome-specific primary sources. While the current graph facilitates interactive exploration and basic analyses, its limited node attribute density poses challenges for advanced machine learning applications following GNN embedding. To address this constraint, future exploration of alternative graph topologies, including the proposed complex topology in **Figure 9**, is recommended. This study underscores both the synergistic potential of integrating diverse data types and the critical need for expanded data availability to enhance predictive capabilities. Broader testing and reporting in publicly available databases will therefore be pivotal in advancing the value of knowledge graphs in personalized oncology applications.

## Materials and Methods

### Data collection from primary sources

Cancer-specific data was collected from the SQL implementation of the Genomic Data Commons (GDC) database version 22.0 publicly available^24,49^. This included cancer patient IDs (*case_id_id*) linked to their tumor primary site where the cancer started developing. Through the manuscript we refer to primary sites as cancer types for simplicity. Somatic mutations leading to amino acid substitutions were also collected from this dataset and linked to the patient in which they occurred. Finally, results from the differential expression analysis in cancer *vs*. normal samples conducted per cancer type in this dataset were also extracted for all available genes. All GDC-derived data was extracted with HUGO Gene Nomenclature Committee (HGNC) gene symbols. For reference, mutations were also annotated based on their inclusion in the natural variance dataset 1000 Genomes^24^.

A PPI phosphorylation network of the kinome was collected from the work of Olow *et al*.^42^ Nodes in the PPI network represent protein coding genes of kinases and their substrates, identified by their HGNC symbols. The network was reconstructed using the edges file, and the nodes file was used to get node attributes, in particular the node subtype (“kinase”, “substrate”, or “both”). Nodes with subtype “both” were annotated as “kinase”.

Bioactivity differences between mutants and wild-type proteins were computed as previously described for the dataset of mutants annotated in ChEMBL 31 and the Papyrus dataset (version 5.5)^50^. In particular, the Wasserstein distance was calculated between the bioactivity distributions of all annotated mutants and the wild-type protein. ChEMBL 31 was also used to extract drugs in all phases of development (0: preclinical, 1-3: clinical, 4: approved) linked via their mechanism of action to particular proteins and mutations. All proteins were represented by their UniProt accession codes.

The structural location of mutations was assessed twofold. The first approach was protein family-independent and entailed the calculation of the average distance between the mutated residues’ centroid and the co-crystallized ligand’s centroid across available structures for the protein in the PDB. All proteins were represented by their UniProt accession codes. This method was previously detailed^50^. The second approach was kinome-specific and entailed querying the KLIFS database.^45^ Through the API, the protein-coding gene’s HGNC symbol was linked to all available structures in the database. These structures were used to query the 85-residue KLIFS binding pocket and the residues forming it. Finally, a consensus KLIFS binding pocket was defined for each queried gene by selecting the most representative residue numbers for each aligned position in the pocket.

Amino acid substitutions were annotated with their corresponding Epstein coefficient of difference^65^ and Grantham’s distance^66^, as previously defined^50^. The Epstein coefficient of difference is directional and was therefore annotated for each individual substitution. Grantham’s distance, on the other hand, is non-directional and was therefore linked to each absolute amino acid change independently of its direction. The Blosum62 score^67^ was also included as a metric to define the likelihood of an amino acid substitution to happen more or less frequently than random change, based on evolutionary conservation.

### Ontology mapping and protein family annotation

The complete dataset from The Human Protein Atlas resource^81^ was downloaded to facilitate protein family filtering and ontology mapping between proteins (UniProt accession codes) and genes (HGNC gene symbols). The downloaded tab-separated file contained a subset of the data from version 23.0 of the resource corresponding to the fields available in the data portal. Kinases were annotated by selecting entries containing the term “Kinase” in their *Molecular function* field. Receptor tyrosine kinases (RTKs) were annotated as kinases additionally containing the term “Receptor” in their *Molecular function* field. Membrane proteins were annotated as those containing the term “Plasma membrane” in their

*Subcellular main location* field. Substrates in the PPI phosphorylation network were further annotated as members of two membrane protein families of interest, G protein-coupled receptors (GPCRs) and solute carriers (SLCs). The former were filtered when the field *Protein class* contained the term “G-protein coupled”. The latter when the field *Gene* started with “SLC”.

### Graph building and interactive exploration

The knowledge graph constructed contained gene and mutation nodes. Gene nodes were all nodes from the PPI phosphorylation network, including both kinases and substrates, and kinases with somatic mutations in the GDC dataset. Edges between gene nodes were directly derived from the PPI network. Mutation nodes were kinase somatic mutations in the GDC dataset and kinase mutations with bioactivity annotations obtained from the dataset previously constructed^50^. Edges between mutations represented co-occurrence in the same patient of the GDC dataset. Mutation and gene nodes were connected by edges representing the association between mutations happening in a specific gene. Cancer mutations were additionally linked to their gene with an edge representing the patient where the mutation occurs. Gene nodes (both kinases and substrates) were annotated with differential expression data from GDC for all available cancer types. Mutation nodes were annotated with bioactivity and structural data when available. Structural data was annotated based on the mutation residue. All mutations were annotated with the Epstein coefficient of difference, based on the amino acid substitution. All nodes (genes and mutations) were annotated with the number of drugs in different phases of development that were associated to the protein encoded by the gene or the mutation in their mechanism of action. Gene nodes were further annotated as part of certain protein families of interest (membrane proteins, RTKs, GPCRs, SLCs). These data were saved in nodes and edges files.

NetworkX^53^ was used to build a multigraph in python. The graph was built from the edges file and the nodes were annotated with their attributes using the nodes file. A package was constructed in python to enable modular build, storage, and updates of the graph. To this end, a graph metadata file is created every time a graph is initialized with a new combination of graph name and edges/nodes files. The package also contains modules to facilitate visual interactive exploration of the data in the knowledge graph based on the graph visualization python package IPysigma^77^.

### Network analysis

Network analysis algorithms implemented in NetworkX were used to explore the knowledge graph and pinpoint relevant nodes. The network node metrics calculated were degree, degree centrality, betweenness centrality, closeness centrality, eigenvector centrality, and information centrality. The python package built for this project enabled the calculation of these metrics for the complete network but also for subsets of it. Precomputed subsets included those containing only one type of edge: cancer co-occurrence, phosphorylation, and gene-mutation association. This distinction enabled the identification of key nodes in each of those layers of information. Additionally, each cancer type was analyzed independently by computing network subsets containing only edges corresponding to patients in a specific cancer type and the phosphorylation edges linking the subset genes. The package analysis modules not only enable the separate examination of these and other subsets, but also facilitate the integration of the subset analysis results into the entire knowledge graph. The computed metrics can be further explored interactively.

## Data availability

The code and data underlying the results and conclusions derived from this work are available online at Zenodo (https://doi.org/10.5281/zenodo.11236776). This repository also contains the kinome and RTK graph files and RTK interactive visualization sessions.

## References

1. Naga, D., Muster, W., Musvasva, E. & Ecker, G. F. Off-targetP ML: an open source machine learning framework for off-target panel safety assessment of small molecules. Journal of Cheminformatics 14, 27 (2022).

2. Kalsekar, I., Koehler, J. & Mulvaney, J. Impact of ACE inhibitors on mortality and morbidity in patients with AMI: Does tissue selectivity matter? Value in Health 14, 184–191 (2011).

3. Montoya, S. et al. Targeted Therapies in Cancer: To Be or Not to Be, Selective. Biomedicines 9, 1591 (2021).

4. Vlot, A. H. C. et al. Target and Tissue Selectivity Prediction by Integrated Mechanistic Pharmacokinetic-Target Binding and Quantitative Structure Activity Modeling. AAPS J 20, 11 (2017).

5. Blagosklonny, M. V. Selective protection of normal cells from chemotherapy, while killing drug-resistant cancer cells. Oncotarget 14, 193–206 (2023).

6. Zhao, Z., Ukidve, A., Kim, J. & Mitragotri, S. Targeting Strategies for Tissue-Specific Drug Delivery. Cell 181, 151– 167 (2020).

7. Lassen, U. N. et al. Precision oncology: a clinical and patient perspective. Future Oncology 17, 3995–4009 (2021).

8. Pessoa, J., Martins, M., Casimiro, S., Pérez-Plasencia, C. & Shoshan-Barmatz, V. Editorial: Altered Expression of Proteins in Cancer: Function and Potential Therapeutic Targets. Front Oncol 12, 949139 (2022).

9. Waarts, M. R., Stonestrom, A. J., Park, Y. C. & Levine, R. L. Targeting mutations in cancer. J Clin Invest 132, e154943 (2022).

10. Bhullar, K. S. et al. Kinase-targeted cancer therapies: progress, challenges and future directions. Molecular Cancer 17, 48 (2018).

11. Liu, G., Chen, T., Zhang, X., Ma, X. & Shi, H. Small molecule inhibitors targeting the cancers. MedComm 3, e181 (2022).

12. Liu, X. et al. Cryo-EM structures of cancer-specific helical and kinase domain mutations of PI3Kα. Proceedings of the National Academy of Sciences 119, e2215621119 (2022).

13. Dixit, A. et al. Sequence and Structure Signatures of Cancer Mutation Hotspots in Protein Kinases. PLOS ONE 4, e7485 (2009).

14. Kong, Y. et al. Small Molecule Inhibitors as Therapeutic Agents Targeting Oncogenic Fusion Proteins: Current Status and Clinical. Molecules 28, 4672 (2023).

15. Miller, M. S. et al. Identification of allosteric binding sites for PI3Kα oncogenic mutant specific inhibitor design. Bioorganic & Medicinal Chemistry 25, 1481–1486 (2017).

16. Kim, P., Zhao, J., Lu, P. & Zhao, Z. mutLBSgeneDB: mutated ligand binding site gene DataBase. Nucleic Acids Res 45, D256–D263 (2017).

17. Zubair, T. & Bandyopadhyay, D. Small Molecule EGFR Inhibitors as Anti-Cancer Agents: Discovery, Mechanisms of Action, and Opportunities. International Journal of Molecular Sciences 24, 2651 (2023).

18. Varkaris, A. et al. Allosteric PI3Kα Inhibition Overcomes On-target Resistance to Orthosteric Inhibitors Mediated by Secondary PIK3CA Mutations. Cancer Discovery 14, 227–239 (2024).

19. Dupont, C. A., Riegel, K., Pompaiah, M., Juhl, H. & Rajalingam, K. Druggable genome and precision medicine in cancer: current challenges. The FEBS Journal 288, 6142–6158 (2021).

20. Radoux, C. J., Vianello, F., McGreig, J., Desai, N. & Bradley, A. R. The druggable genome: Twenty years later. Front. Bioinform. 2, 958378 (2022).

21. Gorostiola González, M., Janssen, A. P. A., IJzerman, A. P., Heitman, L. H. & van Westen, G. J. P. Oncological drug discovery: AI meets structure-based computational research. Drug Discovery Today 27, 1661–1670 (2022).

22. Pacini, C. et al. A comprehensive clinically informed map of dependencies in cancer cells and framework for target prioritization. Cancer Cell 42, 301–316 (2024).

23. Singha, M. et al. Unlocking the Potential of Kinase Targets in Cancer: Insights from CancerOmicsNet, an AI-Driven Approach to Drug Response Prediction in Cancer. Cancers (Basel) 15, 4050 (2023).

24. Bongers, B. J. et al. Pan-cancer functional analysis of somatic mutations in G protein-coupled receptors. Scientific Reports 12, 21534 (2022).

25. Zhao, J., Cheng, F., Wang, Y., Arteaga, C. L. & Zhao, Z. Systematic Prioritization of Druggable Mutations in ∼5000 Genomes Across 16 Cancer Types Using a Structural Genomics-based Approach. Molecular & Cellular Proteomics 15, 642–656 (2016).

26. Silva, M. C., Eugénio, P., Faria, D. & Pesquita, C. Ontologies and Knowledge Graphs in Oncology Research. Cancers (Basel) 14, 1906 (2022).

27. Zhang, W., Chien, J., Yong, J. & Kuang, R. Network-based machine learning and graph theory algorithms for precision oncology. npj Precision Oncology 1, 25 (2017).

28. Cesareni, G., Sacco, F. & Perfetto, L. Assembling Disease Networks From Causal Interaction Resources. Front. Genet. 12, 694468 (2021).

29. Gogleva, A. et al. Knowledge graph-based recommendation framework identifies drivers of resistance in EGFR mutant non-small cell lung cancer. Nat Commun 13, 1667 (2022).

30. Wang, Y. & Liu, Z.-P. Identifying biomarkers for breast cancer by gene regulatory network rewiring. BMC Bioinformatics 22, 308 (2022).

31. Zhao, L. et al. Biological knowledge graph-guided investigation of immune therapy response in cancer with graph neural network. Briefings in Bioinformatics 24, bbad023 (2023).

32. Pu, L. et al. An integrated network representation of multiple cancer-specific data for graph-based machine learning. npj Syst Biol Appl 8, 1–8 (2022).

33. Niu, R., Guo, Y. & Shang, X. GLIMS: A two-stage gradual-learning method for cancer genes prediction using multiomics data and co-splicing network. iScience 27, 109387 (2024).

34. Hatano, N., Kamada, M., Kojima, R. & Okuno, Y. Network-based prediction approach for cancer-specific driver missense mutations using a graph neural network. BMC Bioinformatics 24, 383 (2023).

35. Li, B. & Nabavi, S. A multimodal graph neural network framework for cancer molecular subtype classification. BMC Bioinformatics 25, 27 (2024).

36. Quan, X., Cai, W., Xi, C., Wang, C. & Yan, L. AIMedGraph: a comprehensive multi-relational knowledge graph for precision medicine. Database 2023, baad006 (2023).

37. Bang, D., Lim, S., Lee, S. & Kim, S. Biomedical knowledge graph learning for drug repurposing by extending guilt-by-association to multiple layers. Nat Commun 14, 3570 (2023).

38. Renaux, A. et al. A knowledge graph approach to predict and interpret disease-causing gene interactions. BMC Bioinformatics 24, 324 (2023).

39. Maghawry, N., Ghoniemy, S., Shaaban, E. & Emara, K. An Automatic Generation of Heterogeneous Knowledge Graph for Global Disease Support: A Demonstration of a Cancer Use Case. Big Data and Cognitive Computing 7, 21 (2023).

40. Chandak, P., Huang, K. & Zitnik, M. Building a knowledge graph to enable precision medicine. Sci Data 10, 67 (2023).

41. Jin, S., Liang, H., Zhang, W. & Li, H. Knowledge Graph for Breast Cancer Prevention and Treatment: Literature-Based Data Analysis Study. JMIR Medical Informatics 12, e52210 (2024).

42. Olow, A. et al. An atlas of the human kinome reveals the mutational landscape underlying dysregulated phosphorylation cascades in cancer. Cancer Research 76, 1733–1745 (2016).

43. Jensen, M. A., Ferretti, V., Grossman, R. L. & Staudt, L. M. The NCI Genomic Data Commons as an engine for precision medicine. Blood 130, 453–459 (2017).

44. Burley, S. K. et al. RCSB Protein Data Bank: biological macromolecular structures enabling research and education in fundamental biology, biomedicine, biotechnology and energy. Nucleic Acids Res 47, D464–D474 (2019).

45. Kanev, G. K., de Graaf, C., Westerman, B. A., de Esch, I. J. P. & Kooistra, A. J. KLIFS: an overhaul after the first 5 years of supporting kinase research. Nucleic Acids Res 49, D562–D569 (2021).

46. Mendez, D. et al. ChEMBL: towards direct deposition of bioassay data. Nucleic Acids Res. 47, D930–D940 (2019).

47. Danesi, R. et al. Druggable targets meet oncogenic drivers: opportunities and limitations of target-based classification of tumors and the role of Molecular Tumor Boards. ESMO Open 6, 100040 (2021).

48. Yang, Y., Li, S., Wang, Y., Zhao, Y. & Li, Q. Protein tyrosine kinase inhibitor resistance in malignant tumors: molecular mechanisms and future perspective. Sig Transduct Target Ther 7, 1–36 (2022).

49. Bongers, B. et al. Data underlying the article: Pan-cancer in silico analysis of somatic mutations in G-protein coupled receptors: The effect of evolutionary conservation and natural variance. Available at 10.4121/15022410.V1 (2021).

50. Gorostiola González, M. et al. Excuse me, there is a mutant in my bioactivity soup! A comprehensive analysis of the genetic variability landscape of bioactivity databases and its effect on activity modelling. Preprint at ChemRxiv 10.26434/chemrxiv-2024-kxlgm (2024).

51. Geleta, D. et al. Biological Insights Knowledge Graph: an integrated knowledge graph to support drug development. 2021.10.28.466262 Preprint at ArXiv 10.1101/2021.10.28.466262 (2021).

52. Rozemberczki, B. et al. MOOMIN: Deep Molecular Omics Network for Anti-Cancer Drug Combination Therapy. Preprint at ArXiv 10.48550/arXiv.2110.15087 (2022).

53. Hagberg, A. A., Schult, D. A. & Swart, P. J. Exploring Network Structure, Dynamics, and Function using NetworkX. in Proceedings of the 7th Python in Science Conference (SciPy 2008) (2008).

54. Karim, M. R. et al. From Large Language Models to Knowledge Graphs for Biomarker Discovery in Cancer. Preprint at ArXiv 10.48550/arXiv.2310.08365 (2023).

55. Pan, J. Z. et al. Large Language Models and Knowledge Graphs: Opportunities and Challenges. Preprint at ArXiv http://arxiv.org/abs/2308.06374 (2023).

56. Zhang, N. et al. Predicting Anticancer Drug Responses Using a Dual-Layer Integrated Cell Line-Drug Network Model. PLOS Computational Biology 11, e1004498 (2015).

57. Yang, Z. et al. A mutation-induced drug resistance database (MdrDB). Commun Chem 6, 1–9 (2023).

58. Sha, D. et al. Tumor Mutational Burden (TMB) as a Predictive Biomarker in Solid Tumors. Cancer Discov 10, 1808– 1825 (2020).

59. Oh, J.-H. et al. Spontaneous mutations in the single TTN gene represent high tumor mutation burden. npj Genom. Med. 5, 1–11 (2020).

60. Ibáñez, M. et al. The Mutational Landscape of Acute Promyelocytic Leukemia Reveals an Interacting Network of Co-Occurrences and Recurrent Mutations. PLOS ONE 11, e0148346 (2016).

61. Zhong, X., Yang, H., Zhao, S., Shyr, Y. & Li, B. Network-based stratification analysis of 13 major cancer types using mutations in panels of cancer genes. BMC Genomics 16, S7 (2015).

62. Li, B., Wang, T. & Nabavi, S. Cancer molecular subtype classification by graph convolutional networks on multiomics data. in Proceedings of the 12th ACM Conference on Bioinformatics, Computational Biology, and Health Informatics 1–9 (2021).

63. Guo, H., Lv, X., Li, Y. & Li, M. Attention-based GCN integrates multi-omics data for breast cancer subtype classification and patient-specific gene marker identification. Brief Funct Genomics 22, 463–474 (2023).

64. Tanvir, R. B., Islam, M. M., Sobhan, M., Luo, D. & Mondal, A. M. MOGAT: A Multi-Omics Integration Framework Using Graph Attention Networks for Cancer Subtype Prediction. International Journal of Molecular Sciences 25, 2788 (2024).

65. Epstein, C. J. Non-randomness of Ammo-acid Changes in the Evolution of Homologous Proteins. Nature 215, 355– 359 (1967).

66. Grantham, R. Amino acid difference formula to help explain protein evolution. Science 185, 862–864 (1974).

67. Henikoff, S. & Henikoff, J. G. Amino acid substitution matrices from protein blocks. Proc Natl Acad Sci U S A 89, 10915–10919 (1992).

68. Martínez-Sáez, O. et al. Frequency and spectrum of PIK3CA somatic mutations in breast cancer. Breast Cancer Research 22, 45 (2020).

69. Boyer, S., Money-Kyrle, S. & Bent, O. Predicting protein stability changes under multiple amino acid substitutions using equivariant graph neural networks. Preprint at ArXiv http://arxiv.org/abs/2305.19801 (2023).

70. Robichaux, J. P. et al. Structure-based classification predicts drug response in EGFR-mutant NSCLC. Nature 597, 732–737 (2021).

71. Van Linden, O. P. J., Kooistra, A. J., Leurs, R., De Esch, I. J. P. & De Graaf, C. KLIFS: A Knowledge-Based Structural Database To Navigate Kinase− Ligand Interaction Space. Journal of Medicinal Chemistry 57, 249–277 (2013).

72. Engin, H. B., Kreisberg, J. F. & Carter, H. Structure-Based Analysis Reveals Cancer Missense Mutations Target Protein Interaction Interfaces. PLOS ONE 11, e0152929 (2016).

73. Pujara, J., Augustine, E. & Getoor, L. Sparsity and Noise: Where Knowledge Graph Embeddings Fall Short. In Proceedings of the 2017 Conference on Empirical Methods in Natural Language Processing 1751–1756 (2017).

74. Xu, J., Zhang, W., Duan, Q. & Li, S. HOAP: Node attribute completion of knowledge graph based on high-order neighbor attribute propagation. Preprint at ArXiv 10.21203/rs.3.rs-1937079/v1 (2022).

75. Guo, D., Chu, Z. & Li, S. Fair Attribute Completion on Graph with Missing Attributes. Preprint at ArXiv http://arxiv.org/abs/2302.12977 (2023).

76. Gorostiola González, M. et al. Computational Characterization of Membrane Proteins as Anticancer Targets: Current Challenges and Opportunities. International Journal of Molecular Sciences 25, 3698 (2024).

77. Plique, G. ipysigma. Zenodo 10.5281/zenodo.7521476 (2023).

78. Boutros, A. et al. Activity and safety of first-line treatments for advanced melanoma: A network meta-analysis. European Journal of Cancer 188, 64–79 (2023).

79. Bosdriesz, E. et al. Identifying mutant-specific multi-drug combinations using comparative network reconstruction. iScience 25, 104760 (2022).

80. Hu, C. et al. Optimizing drug combination and mechanism analysis based on risk pathway crosstalk in pan cancer. Sci Data 11, 74 (2024).

81. Uhlén, M. et al. Tissue-based map of the human proteome. Science 347, 1260419 (2015).

