## Supplementary Tables 1-6 and Figures 1-11 for "A patient-centric knowledge graph approach to prioritize mutants for selective anti-cancer targeting"

**Supplementary Table 1.** Top 25 cancer mutation nodes with the highest mutation frequency (pan-cancer) in the kinome knowledge graph.

| **Cancer mutation** | **Frequency** |
|---|---|
| BRAF_V600E | 565 |
| PIK3CA_E545K | 258 |
| PIK3CA_H1047R | 234 |
| PIK3CA_E542K | 167 |
| PIK3CA_R88Q | 68 |
| AKT1_E17K | 53 |
| BRAF_V600M | 40 |
| FGFR3_S249C | 39 |
| ERBB2_S310F | 38 |
| PIK3CA_H1047L | 37 |
| PIK3CA_N345K | 34 |
| PIK3CA_E726K | 30 |
| PIK3CA_G118D | 28 |
| FGFR2_S252W | 26 |
| ERBB3_V104M | 25 |
| EGFR_L858R | 23 |
| PIK3CA_C420R | 23 |
| PIK3CA_Q546R | 22 |
| EGFR_A289V | 21 |
| PIK3CA_E453K | 19 |
| EGFR_G598V | 19 |
| PIK3CA_R108H | 19 |
| PIK3CA_E545A | 18 |
| PIK3CA_M1043I | 18 |
| MAPK1_E322K | 18 |

**Supplementary Table 2.** Top 25 ranked cancer mutation nodes in the kinome knowledge graph according to their degree. For reference, the mutation frequency across all cancer patients analyzed is reported, and the rank that would correspond to the cancer mutation if calculated based on the mutation frequency, if this rank is 1-25 (otherwise reported as >25). Additionally, the number of unique mutations reported for the gene is reported. *Mutations present in the natural variance dataset 1000 Genomes.

| **Cancer mutation** | **Degree** | **Rank** | **Mutation frequency** | **Frequency rank** | **Gene cancer mutations** |
|---|---|---|---|---|---|
| PIK3CA_R88Q | 10275 | 1 | 68 | 5 | 268 |
| BRAF_V600E | 6219 | 2 | 565 | 1 | 132 |
| PRKDC_R2522Q | 3376 | 3 | 11 | >25 | 521 |
| MTOR_R2152C | 3042 | 4 | 4 | >25 | 345 |
| NEK3_S284L* | 2898 | 5 | 6 | >25 | 52 |
| PAK5_E144K | 2705 | 6 | 7 | >25 | 203 |
| GCK_A2V | 2661 | 7 | 4 | >25 | 79 |
| PIK3CA_E545K | 2484 | 8 | 258 | 2 | 268 |
| PDK2_A259V | 2483 | 9 | 3 | >25 | 40 |
| PIK3CA_H1047R | 2439 | 10 | 234 | 3 | 268 |
| TTN_R2506Q | 2409 | 11 | 8 | >25 | 7791 |
| CAMK1D_S360L | 2352 | 12 | 6 | >25 | 74 |
| TEK_S599L | 2321 | 13 | 4 | >25 | 185 |
| DCAF1_R855Q | 2262 | 14 | 5 | >25 | 153 |
| MAP3K15_R493W | 2191 | 15 | 7 | >25 | 199 |
| TTN_D19391N | 2175 | 16 | 8 | >25 | 7791 |
| ROCK1_R1012Q | 2159 | 17 | 5 | >25 | 186 |
| SMG1_R803H | 2127 | 18 | 3 | >25 | 386 |
| HIPK1_R875H | 2123 | 19 | 5 | >25 | 148 |
| ROCK1_R590Q | 2118 | 20 | 5 | >25 | 186 |
| STK3_S344L | 2089 | 21 | 5 | >25 | 963 |
| ROCK2_R339Q | 2071 | 22 | 4 | >25 | 153 |
| TTN_R33466C* | 2064 | 23 | 4 | >25 | 7791 |
| DGKB_R685Q* | 2059 | 24 | 3 | >25 | 217 |
| IP6K1_R329H | 2054 | 25 | 4 | >25 | 45 |

**Supplementary Table 3**. Top 25 most frequently mutated proteins per cancer type (as defined by primary site). Mutation frequency is calculated as the sum of all mutations in that protein-cancer type pair.

| **Protein** | **Cancer type** | **Mutation frequency** |
|---|---|---|
| TTN | Skin | 1,931 |
| TTN | Corpus uteri | 1,808 |
| TTN | Bronchus and lung | 1,402 |
| TTN | Colon | 581 |
| TTN | Stomach | 553 |
| PIK3CA | Corpus uteri | 367 |
| PIK3CA | Breast | 357 |
| OBSCN | Corpus uteri | 334 |
| TTN | Bladder | 328 |
| TTN | Brain | 321 |
| BRAF | Thyroid gland | 290 |
| BRAF | Skin | 290 |
| TTN | Breast | 277 |
| TTN | Cervix uteri | 207 |
| TTN | Ovary | 202 |
| OBSCN | Skin | 195 |
| TTN | Rectum | 192 |
| SMG1 | Corpus uteri | 173 |
| LRRK2 | Corpus uteri | 170 |
| TAF1 | Corpus uteri | 168 |
| PRKDC | Corpus uteri | 161 |
| MYO3A | Corpus uteri | 157 |
| OBSCN | Bronchus and lung | 150 |
| OBSCN | Colon | 147 |
| EGFR | Brain | 146 |

**Supplementary Table 4.** Distribution of node and edge types and subtypes across the receptor tyrosine kinase knowledge graph.

| Entity | Type | Subtype | Number of entities |
|---|---|---|---|
| Nodes | Gene | Kinase | 110 |
| | | Receptor kinase (not in PPI as kinase) | 1 |
| | | Substrate | 142 |
| | Mutation | Cancer | 11,660 |
| | | Other (ChEMBL + Papyrus) | 76 |
| Edges | Gene - Gene | Phosphorylation | 673 |
| | Gene - Mutation | Cancer | 13,353 |
| | | Other (ChEMBL + Papyrus) | 76 |
| | Mutation - Mutation | Cancer patient co-occurrence | 127,187 |
| | | Other (ChEMBL + Papyrus multiple substitutions) | 22 |

**Supplementary Table 5.** Top 25 cancer mutation nodes with the highest mutation frequency (pan-cancer) in the receptor tyrosine kinase knowledge graph.

| **Cancer mutation** | **Frequency** |
|---|---|
| FGFR3_S249C | 39 |
| ERBB2_S310F | 38 |
| FGFR2_S252W | 26 |
| ERBB3_V104M | 25 |
| EGFR_L858R | 23 |
| EGFR_A289V | 21 |
| EGFR_G598V | 19 |
| ERBB2_V842I | 17 |
| ERBB2_R678Q | 14 |
| ERBB2_L755S | 13 |
| FGFR2_N550K | 12 |
| FGFR3_Y375C | 10 |
| ERBB2_V777L | 10 |
| EPHA6_R268C | 8 |
| KIT_D816V | 8 |
| FLT3_D835Y | 8 |
| KDR_R1032Q | 8 |
| EGFR_L62R | 7 |
| EGFR_R222C | 7 |
| FGFR2_C383R | 7 |
| MUSK_R854Q | 7 |
| ERBB4_R711C | 7 |
| EGFR_L861Q | 6 |
| KIT_K642E | 6 |
| FGFR1_N577K | 6 |

**Supplementary Table 6.** Top 25 ranked cancer mutation nodes in the receptor tyrosine kinase knowledge graph according to their degree. For reference, the mutation frequency across all cancer patients analyzed is reported, and the rank that would correspond to the cancer mutation if calculated based on the mutation frequency, if this rank is 1-25 (otherwise reported as >25). Additionally, the number of unique mutations reported for the gene is reported. *Mutations present in the natural variance dataset 1000 Genomes.

| **Cancer mutation** | **Degree** | **Rank** | **Mutation frequency** | **Frequency rank** | **Gene cancer mutations** |
|---|---|---|---|---|---|
| TEK_S599L | 391 | 1 | 4 | >25 | 185 |
| EPHA10_D881N | 305 | 2 | 2 | >25 | 140 |
| KDR_R1032Q | 300 | 3 | 8 | 17 | 331 |
| KIT_R888Q | 274 | 4 | 4 | >25 | 214 |
| EPHA6_D243N | 263 | 5 | 6 | >25 | 336 |
| PDGFRA_E156D | 252 | 6 | 2 | >25 | 293 |
| DDR1_D714N | 246 | 7 | 2 | >25 | 112 |
| ERBB3_R916Q* | 237 | 8 | 2 | >25 | 229 |
| FGFR2_R165W | 232 | 9 | 2 | >25 | 166 |
| EPHA4_R745H | 232 | 10 | 2 | >25 | 192 |
| CSF1R_D565N | 229 | 11 | 2 | >25 | 138 |
| INSR_R924Q | 226 | 12 | 2 | >25 | 201 |
| EGFR_R977H | 217 | 13 | 2 | >25 | 265 |
| EPHA6_R788C | 209 | 14 | 2 | >25 | 336 |
| ACVR1C_R245Q | 207 | 15 | 3 | >25 | 91 |
| EPHA2_E523K | 203 | 16 | 2 | >25 | 171 |
| FLT1_E144K* | 201 | 17 | 3 | >25 | 264 |
| PDGFRA_K196N | 200 | 18 | 2 | >25 | 293 |
| EPHA8_A685T | 199 | 19 | 2 | >25 | 185 |
| MET_R412C | 198 | 20 | 2 | >25 | 214 |
| RYK_R563Q | 195 | 21 | 4 | >25 | 64 |
| MET_L982M | 195 | 22 | 2 | >25 | 214 |
| EPHB1_R743W | 194 | 23 | 2 | >25 | 286 |
| EPHA1_R261W | 194 | 24 | 3 | >25 | 133 |
| MUSK_R572K | 192 | 25 | 2 | >25 | 174 |

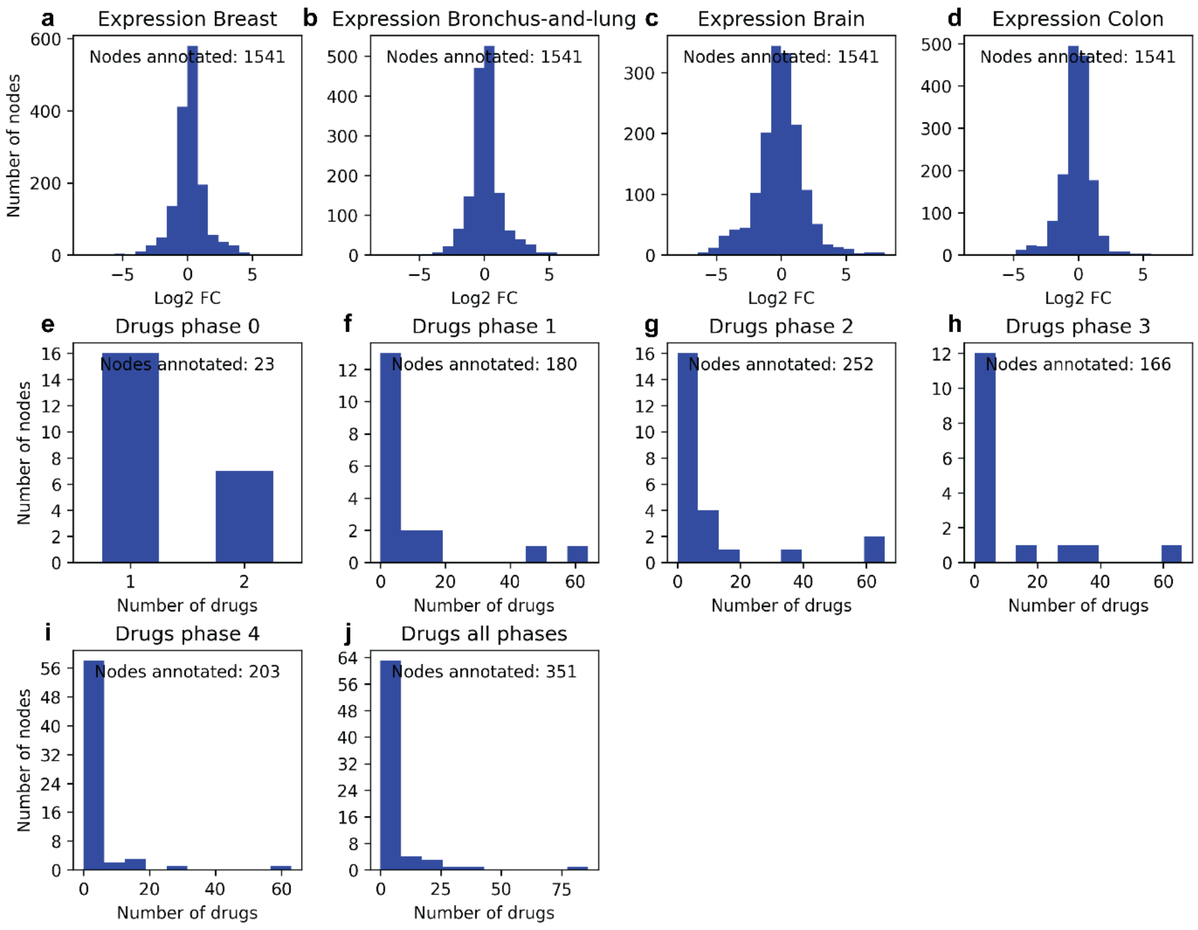


**Supplementary Figure 1.** Density and distribution of gene node attributes. For each of the attributes, the number of gene nodes with non-null values for the particular attribute is depicted on the y-axis. Moreover, the graphs represent the distribution of the attribute values across the gene nodes on the x-axis, in the form of histograms or bar plots, depending on the density of each attribute. Four cancer types are selected as an example to show the distribution of differential expression Log2 fold change (Log2FC) between the tumor tissue and normal tissue (**a-d**). The rest of the graphs represent the number of drugs in different phases of development according to ChEMBL labeling: pre-clinical "0" phase (**e**), clinical phases 1-3 (**f-h**), and approved drugs (**i**). The total number of drugs in any phase is represented in (**j**).

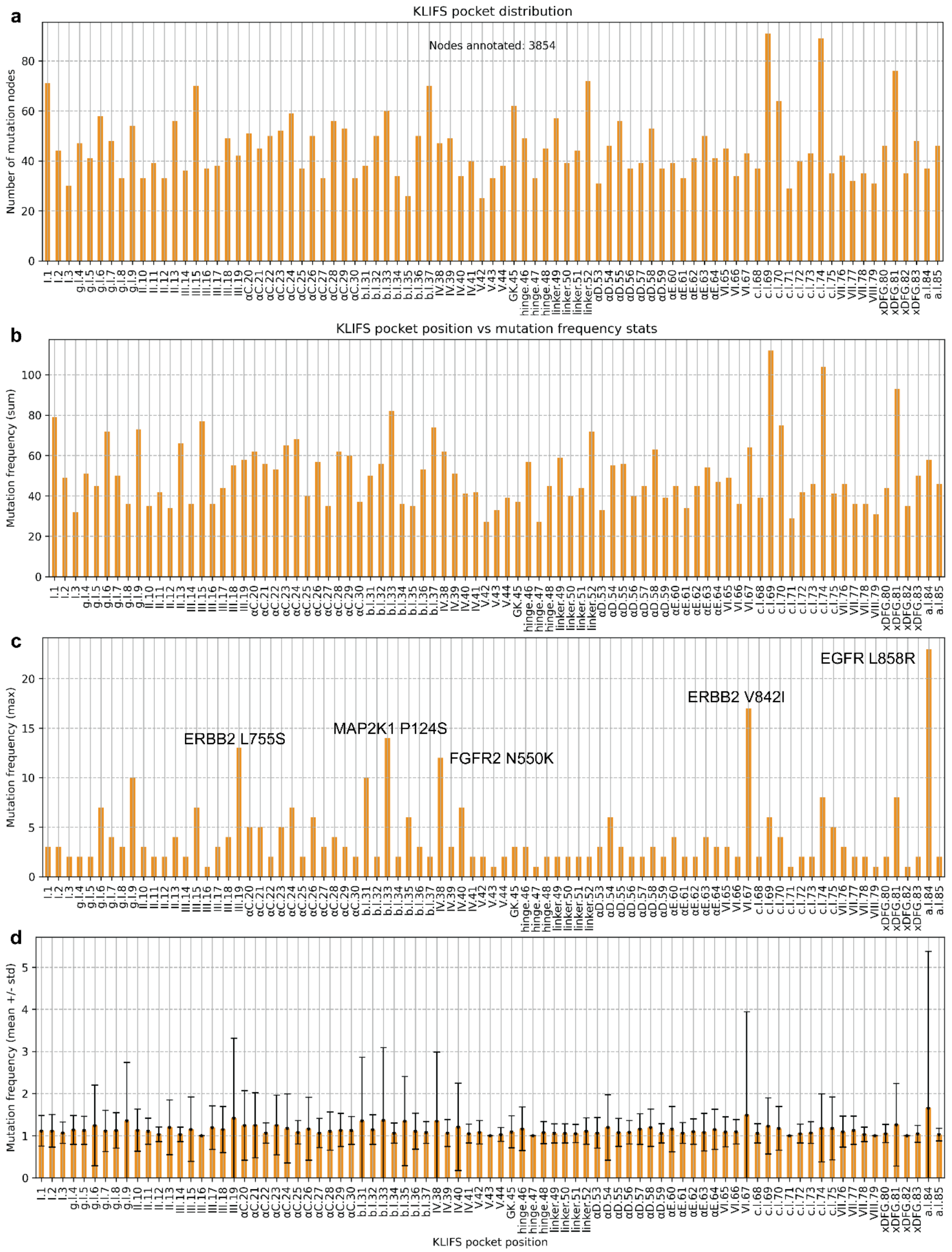


**Supplementary Figure 2.** Density and distribution of KLIFS structural attribute annotations in mutation nodes. **a)** Number of mutation nodes with an annotation for each particular position of the 85-consensus kinase pocket defined by KLIFS. **b-d)** Cancer mutation frequency statistics for each pocket position: sum of mutation frequency in each position (**b**), maximum mutation frequency reported for each position, with the mutations with the top five frequencies labeled (**c**), and mean +/- standard deviation of mutation frequency reported for each pocket position (**d**).

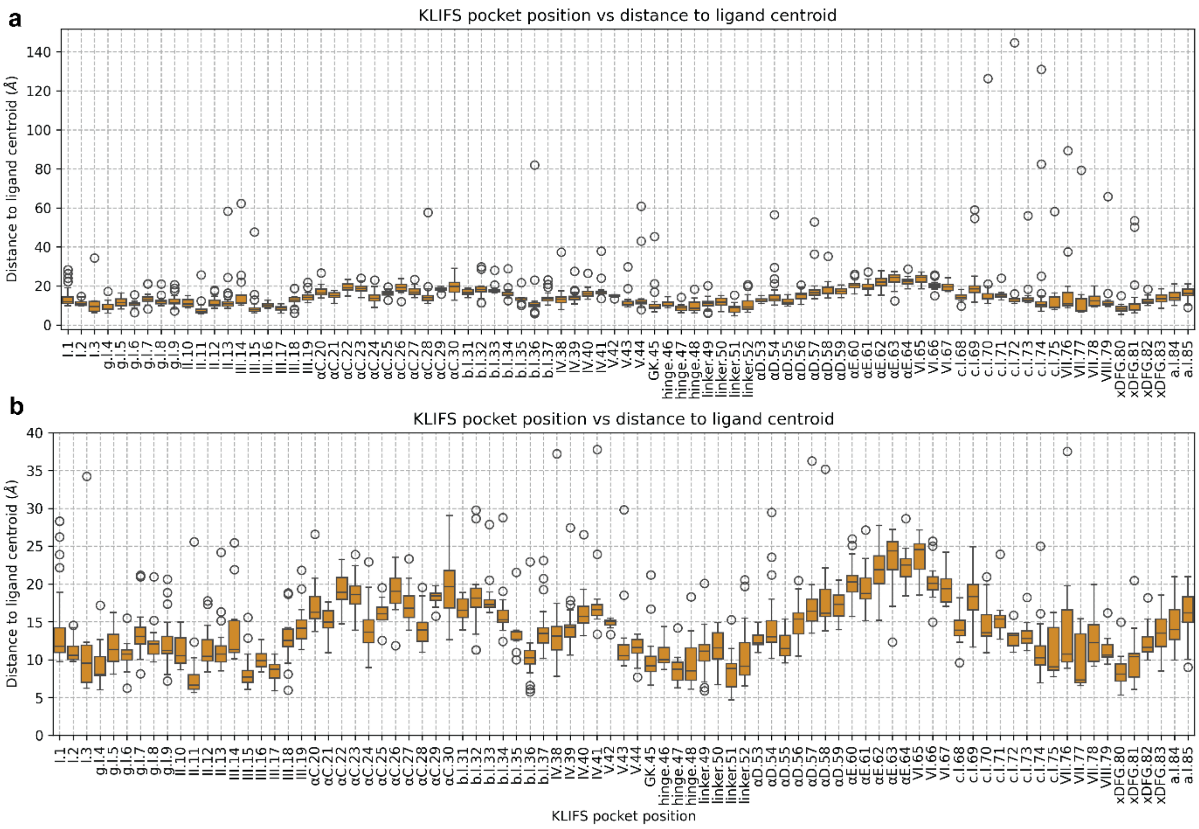


**Supplementary Figure 3.** Average distance to ligand centroid for every KLIFS pocket position calculated for each mutation in that position from available PDB complexes. One distance value is recorded per available mutation in the kinome graph. **a)** Complete distribution**. b)** Distribution of values in the range 0-40 Å.

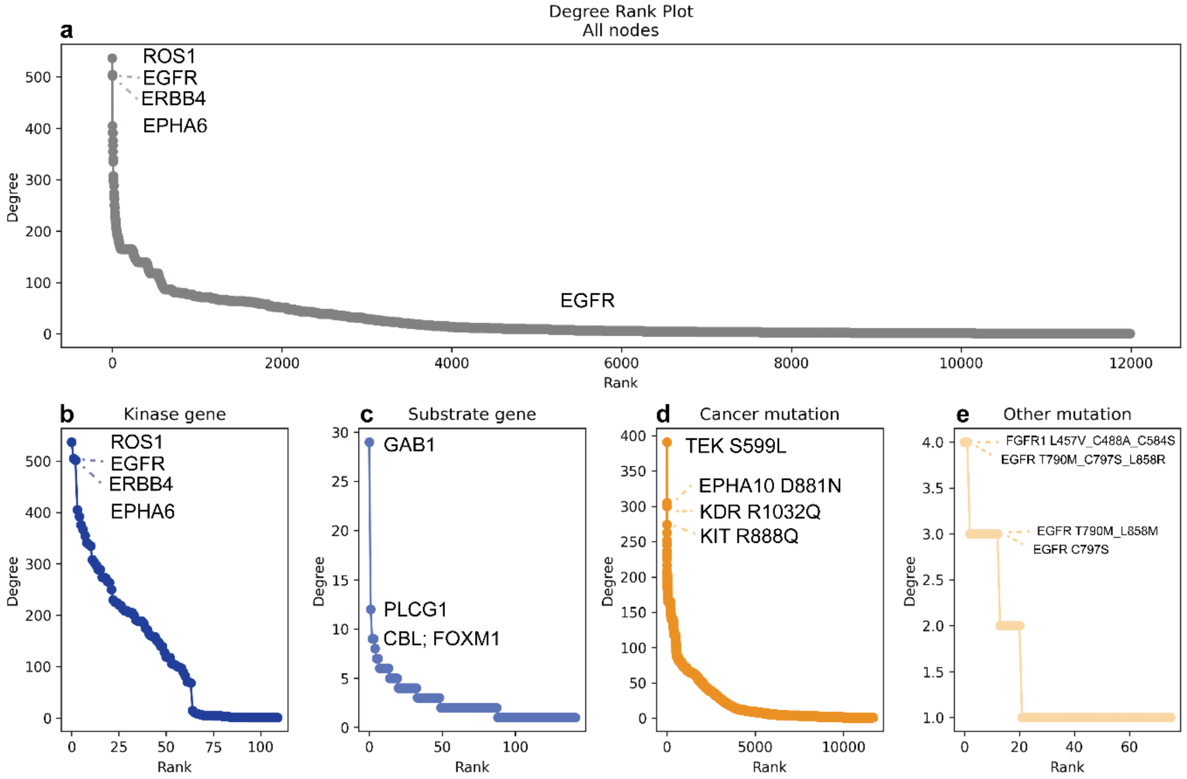


**Supplementary Figure 4.** Node degree rank analysis for the receptor tyrosine kinase knowledge graph. Nodes are ranked based on their degree, which is calculated as the number of edges connecting the node to other nodes in the graph. The degree rank analysis is calculated for all nodes in the graph (**a**) as well as for each node subtype independently: kinase (**b**) and substrate (**c**) gene nodes, and cancer (**d**) and other (**e**) mutations. The top four ranked nodes in each case are labeled accordingly.

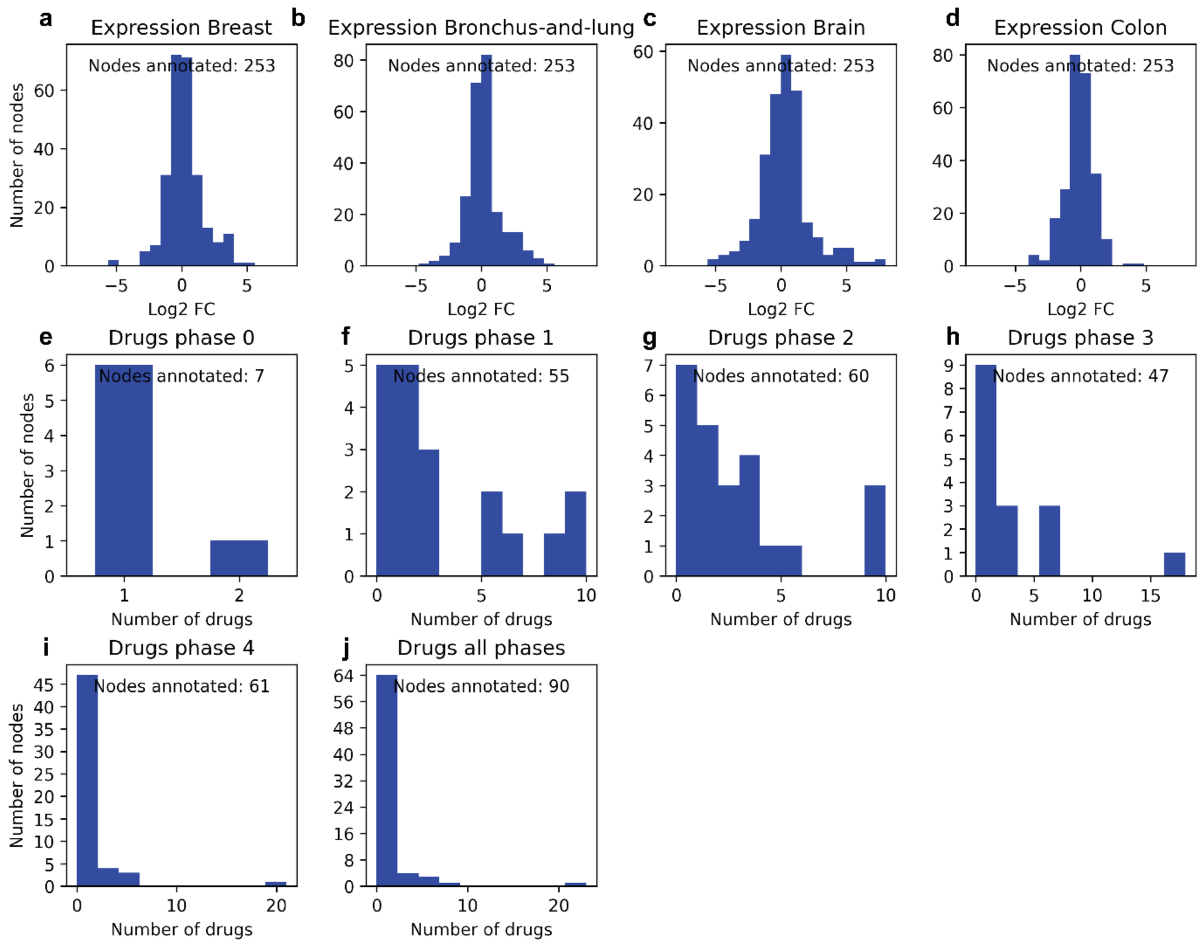


**Supplementary Figure 5.** Density and distribution of gene node attributes in the receptor tyrosine kinase knowledge graph. For each of the attributes, the number of gene nodes with non-null values for the particular attribute is depicted on the y-axis. Moreover, the graphs represent the distribution of the attribute values across the gene nodes on the x-axis, in the form of histograms or bar plots, depending on the density of each attribute. Four cancer types are selected as an example to show the distribution of differential expression Log2 fold change (Log2FC) between the tumor tissue and normal tissue (**a-d**). The rest of the graphs represent the number of drugs in different phases of development according to ChEMBL labeling: pre-clinical “0” phase (**e**), clinical phases 1-3 (**f-h**), and approved drugs (**i**). The total number of drugs in any phase is represented in (**j**).

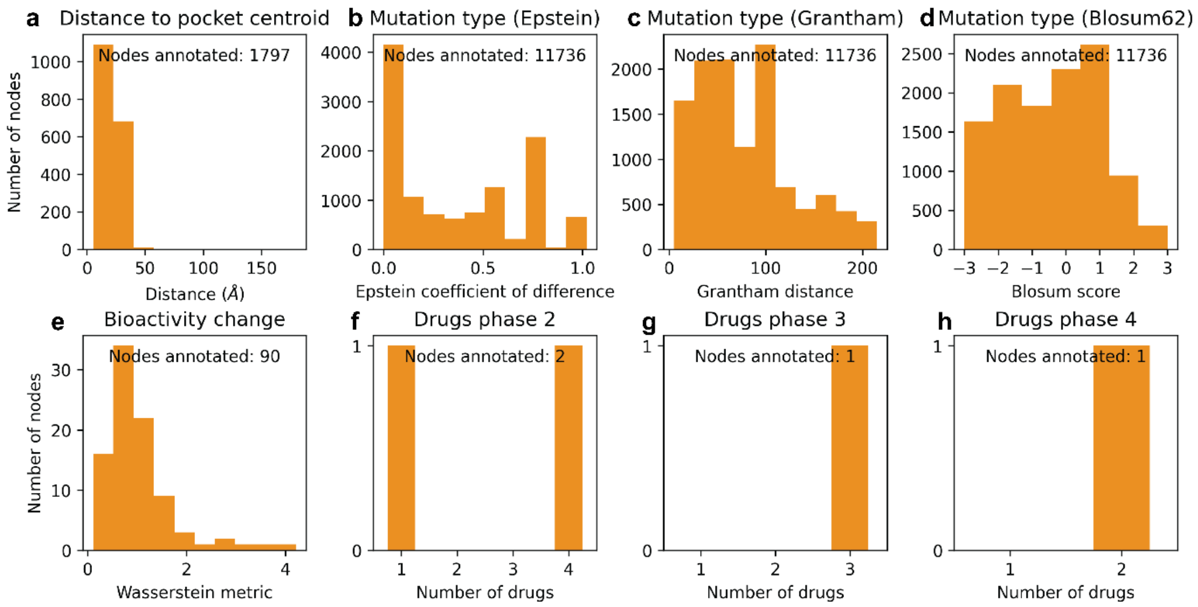


**Supplementary Figure 6**. Density and distribution of mutation node attributes in the receptor tyrosine kinase knowledge graph. For each of the attributes, the number of mutation nodes with non-null values for the particular attribute is depicted on the y-axis. Moreover, the graphs represent the distribution of the attribute values across the mutation nodes on the x-axis, in the form of histograms or bar plots, depending on the density of each attribute. Distribution is represented as histograms for the distance to the pocket centroid calculated from PDB complexes (**a**), the mutation type as determined by the Epstein coefficient of difference (**b**), the mutation type as described by the Grantham distance (**c**), the evolutionary probability of the mutation type as described by the Blosum score in the Blosum62 matrix (**d**), and the bioactivity change represented by the Wasserstein distance between the bioactivity distribution for the mutation and the wild-type protein found in ChEMBL (**e**). Bar plots represent the number of drugs in different phases of development according to ChEMBL labeling: clinical phases 2-3 (**f**-**g**), and approved drugs (**h**). Pre-clinical candidates (phase 0) and drugs in clinical phase 1 are not included because they were not annotated in any mutation nodes.

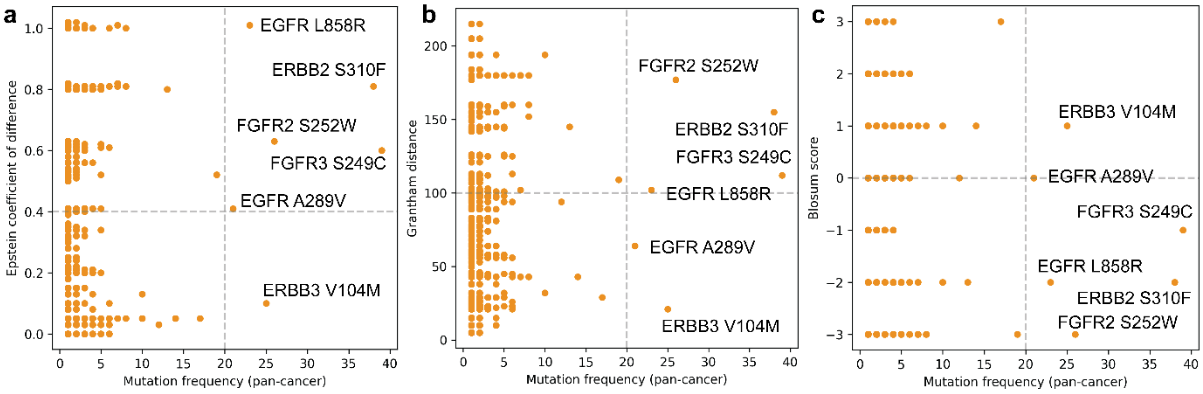


**Supplementary Figure 7.** Correlation between cancer mutation frequency and three metrics describing the amino acid substitution in the kinome knowledge graph: Epstein coefficient of difference (**a**), Grantham distance (**b**), and Blosum score (**c**). Mutations occurring in more than 20 patients pan-cancer are labeled for reference. In **a)**, an Epstein coefficient of difference of 0.4 is taken as an arbitrary threshold to distinguish between conservative (<0.4) and disruptive substitutions (>0.4). In **b)**, a Grantham distance of 100 is taken as an arbitrary threshold to distinguish between conservative (<100) and disruptive substitutions (>100). In **c)**, substitutions with an alignment happening less often than random chance as collected in the Blosum62 matrix are represented by a Blosum score < 0.

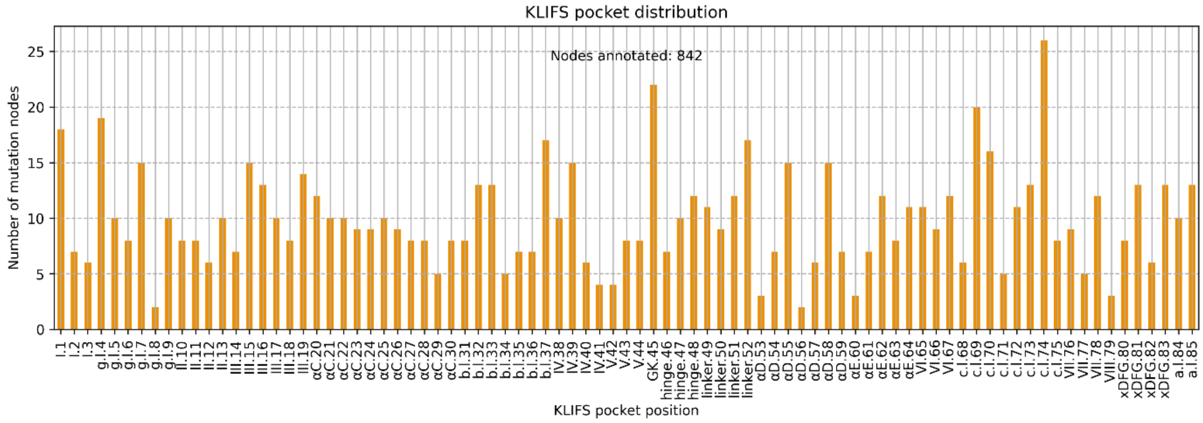


**Supplementary Figure 8.** Number of mutation nodes in the receptor tyrosine kinase graph with an annotation for each particular position of the 85-consensus kinase pocket defined by KLIFS.

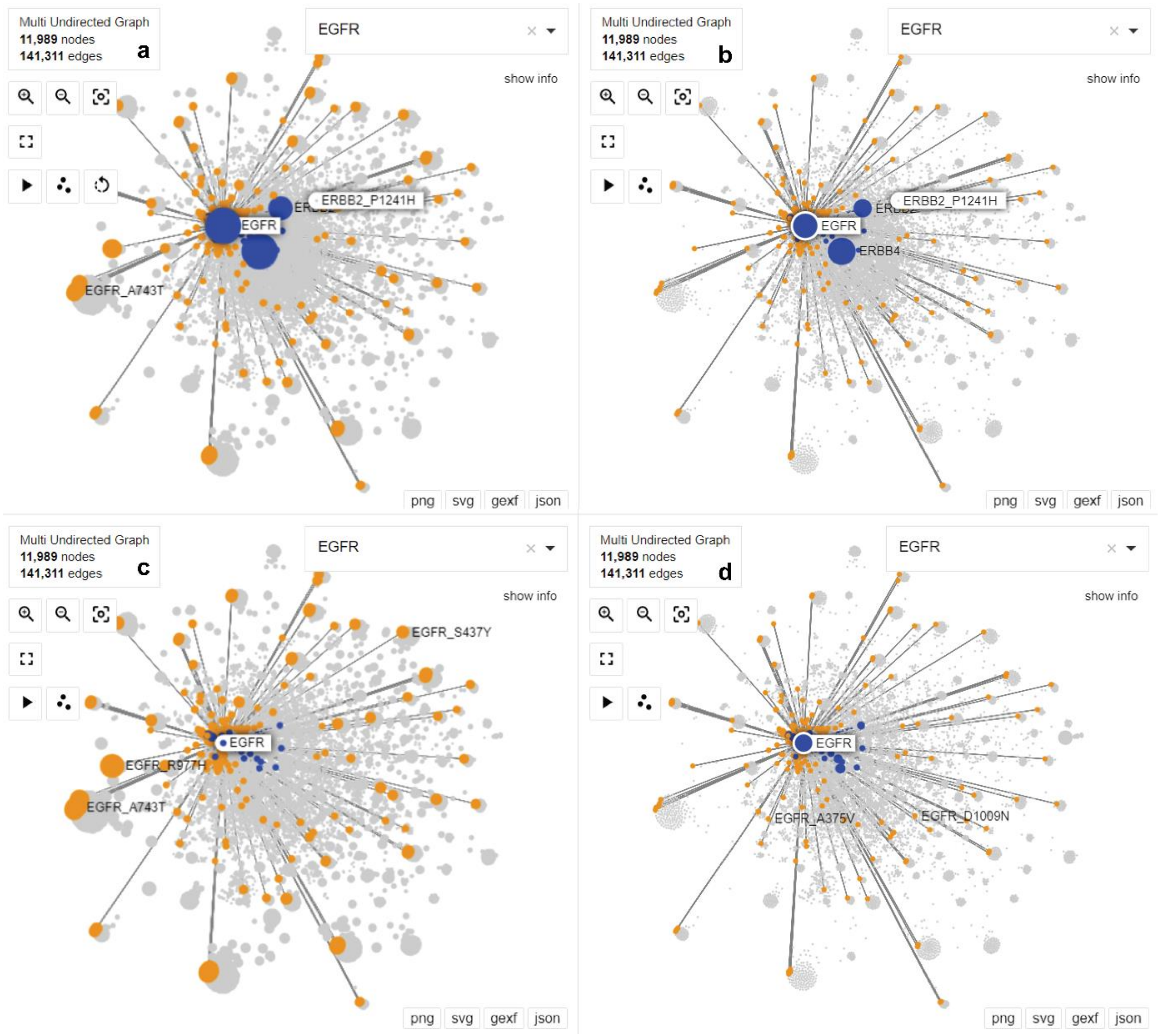


**Supplementary Figure 9.** Node betweenness centrality comparison across layers in the receptor tyrosine kinase knowledge graph, with focus on nodes connected to EGFR for visualization purposes. Gene nodes are represented in blue and mutation nodes are represented in orange. Nodes that are not connected to EGFR are kept in the background and represented in grey. Node size in each panel is determined by the node degree calculated from the whole graph (**a**) or one of the three pre-defined analysis layers: kinase-mutation layer (**b**), cancer-mutation co-occurrence layer (**c**), or phosphorylation layer (**d**).

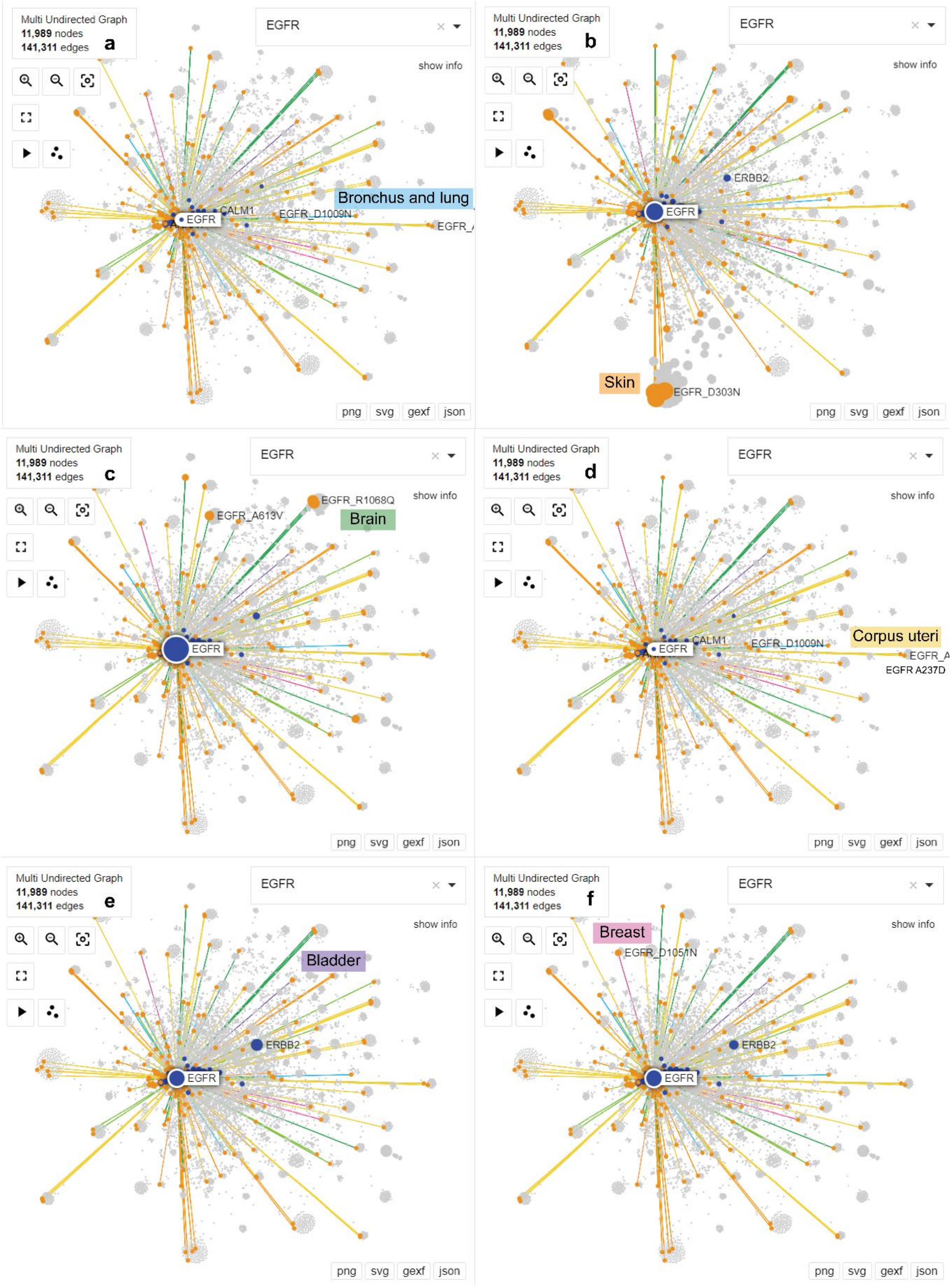


**Supplementary Figure 10.** Node degree comparison across the six most populated cancer types in the receptor tyrosine kinase knowledge graph, with focus on nodes connected to EGFR for visualization purposes. Gene nodes are represented in blue and mutation nodes are represented in orange. Nodes that are not connected to EGFR are kept in the background and represented in grey. Each edge color represents a different cancer type. Node size in each panel is determined by the node degree calculated from the subgraphs for six cancer types: bronchus and lung (**a**, 723 patients – 56 with EGFR mutations. Represented by blue edges), skin (**b**, 356 patients – 29 with EGFR mutations. Represented by orange edges), brain (**c**, 312 patients – 127 with EGFR mutations. Represented by green edges), corpus uteri (**d**, 302 patients – 29 with EGFR mutations. Represented by yellow edges), bladder (**e**, 289 patients – 7 with EGFR mutations. Represented by purple edges), or breast (**f**, 273 patients – 13 with EGFR mutations. Represented by pink edges).

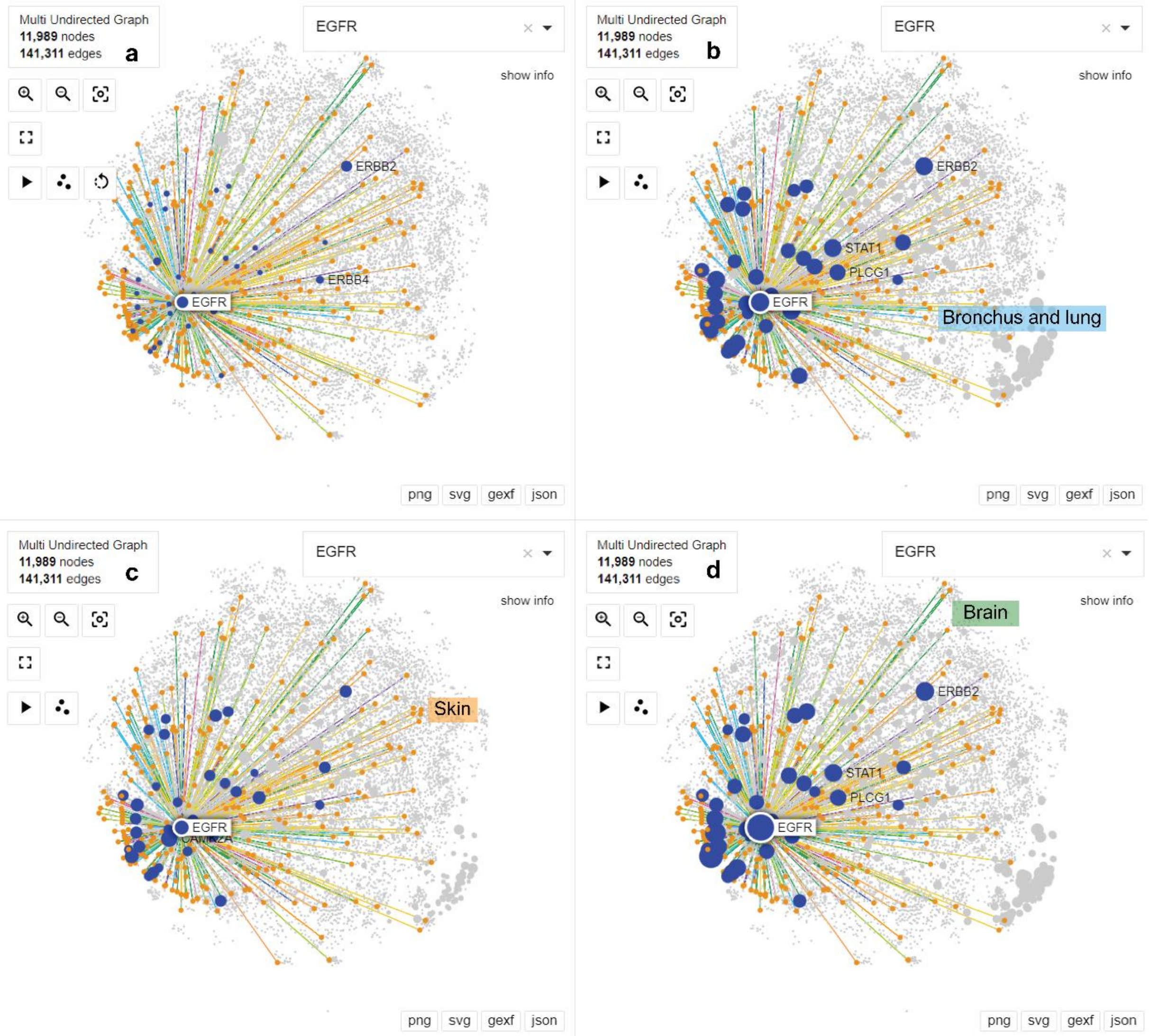


**Supplementary Figure 11.** Comparative visualization of the receptor tyrosine kinase knowledge graph with node sizes representing different attributes. The focus is on nodes connected to EGFR for visualization purposes. Gene nodes are represented in blue and mutation nodes are represented in orange. Nodes that are not connected to EGFR are kept in the background and represented in grey. Each edge color represents a different cancer type. Node size represents the number of approved drugs (**a)** or the differential expression (Log2 fold change) in tumor tissue compared to healthy tissue in the three most populated cancer types: bronchus and lung (**b**, 723 patients – 56 with EGFR mutations. Represented by blue edges), skin (**c**, 356 patients – 29 with EGFR mutations. Represented by orange edges), and brain (**d**, 312 patients – 127 with EGFR mutations. Represented by green edges**).**
